# Dendritic integration in olfactory bulb granule cells: Thresholds for lateral inhibition and role of active conductances upon simultaneous activation

**DOI:** 10.1101/2020.01.10.901397

**Authors:** Max Müller, Veronica Egger

## Abstract

The inhibitory axonless olfactory bulb granule cells (GCs) form reciprocal dendrodendritic synapses with mitral and tufted cells via large spines, mediating recurrent and lateral inhibition. Rat GC dendrites are excitable by local Na^+^ spine spikes and global Ca^2+^- and Na^+^-spikes. To investigate the transition from local to global signaling without Na^+^ channel inactivation we performed simultaneous holographic two-photon uncaging in acute brain slices, along with whole-cell recording and dendritic Ca^2+^ imaging. Less than 10 coactive reciprocal spines were sufficient to generate diverse regional and global signals that also included local dendritic Ca^2+^- and Na^+^-spikes (D-spikes). Individual spines could sense the respective signal transitions as increments in Ca^2+^ entry. Dendritic integration was mostly linear until a few spines below global Na^+^-spike threshold, where often D-spikes set in. NMDARs strongly contributed to active integration, whereas morphological parameters barely mattered. In summary, thresholds for GC-mediated bulbar lateral inhibition are low.

## Introduction

The classical role of dendrites is to receive synaptic or sensory inputs and to conduct the ensuing electrical signals towards the site of action potential (AP) initiation at the axon hillock. While this conduction is passive for smaller membrane depolarizations, recent decades have revealed the presence of active dendritic conductances, most importantly voltage-gated Na^+^ and Ca^2+^ channels (Na_v_s, Ca_v_s) and NMDA receptors (NMDARs), that can amplify locally suprathreshold electrical signals and thus generate dendritic spikes in many neuron types; dendritic Na_v_s also facilitate backpropagation of axonal APs into the dendritic tree (Stuart and Spruston, 2015).

The dendritic integration of multiple excitatory inputs - as detected at the soma - is usually linear for small numbers of coactive synapses but then frequently transitions into regimes of sublinear or supralinear summation with respect to the arithmetic sum of the individual synaptic potentials. The mode of integration depends on dendritic input impedance, the density of active conductances and the distribution of synaptic inputs, both in the spatial and temporal domain (Tran-Van-Minh *et al*., 2015).

For example, supralinear integration occurs in cortical and hippocampal pyramidal cell (PC) dendrites, carried by Ca^2+^-spikes, so-called NMDA-spikes and dendritic Na^+^-spikes (D-spikes) which show as spikelets at the soma (Losonczy and Magee, 2006; Major *et al*., 2013; Larkum *et al*., 2007; Remy *et al*., 2009; Makara and Magee, 2013; Kim *et al*., 2012). Such active integration mechanisms have been shown to e.g. boost axonal AP initiation, enable local coincidence detection and therewith contribute to the induction of synaptic plasticity. Conversely, sublinear integration is performed by e.g. GABAergic cerebellar stellate cell dendrites, that is mainly determined by passive cable properties and reductions in driving force for large dendritic depolarizations (Abrahamsson *et al*., 2012). In contrast to supralinear integration, this type of integration will favor sparse and/or distributed inputs.

Aside from such computations that ultimately convert analogue signals into binary code at the axon initial segment, another functional outcome of dendritic integration is the (possibly graded) release of transmitter from the dendrites themselves. Dendritic transmitter release occurs in many brain regions and is particularly well known from the retina and the olfactory bulb (OB; Ludwig and Pittman, 2003). In the bulb, axonless inhibitory GCs release GABA exclusively from their apical dendrite. These release sites are located within spines that contain reciprocal dendrodendritic synapses with the excitatory mitral and tufted cells (MC/TCs). Unlike principal neurons in the cortex and hippocampus, MC/TCs that belong to different glomerular columns do not communicate directly via axon collaterals or otherwise. Rather, their only interaction happens via lateral inhibition, mediated by GCs and other local interneurons. Although MC/TC-GC synapses are the most numerous in the bulb (Shepherd, 1972; Egger and Urban, 2006), so far it is unknown how many coinciding MC/TC inputs are required to generate global GC activity and with it Ca^2+^ entry also in non-activated spines, possibly invoking lateral inhibition. The properties of dendritic integration in GCs thus critically determine the onset and degree of lateral inhibition.

What is known so far about GC dendritic processing? A single MC/TC input triggers a local Na^+^-AP that is restricted to the spine head because of a high spine neck resistance (Bywalez *et al*., 2015). This spine spike can cause reciprocal release of GABA via gating of high-voltage-activated (HVA) Ca_v_s (Lage-Rupprecht *et al*., 2019). Activation of larger numbers of spines results in global Low-threshold Ca^2+^-spikes (LTS) which are mediated by T-type Ca_v_s (Egger *et al*., 2005; Pressler and Strowbridge, 2019; Pinato and Midtgaard, 2005). Synaptically evoked Na^+^ spikelets have been reported from mouse, turtle and frog GCs, causing regional Ca^2+^ entry (Pinato and Midtgaard, 2005; Zelles *et al*., 2006; Burton and Urban, 2015), while somatic full-blown synaptic Na^+^ APs can be elicited by stimulation of a single glomerulus (Schoppa *et al*., 1998) and result in substantial Ca^2+^ entry throughout the GC dendrite that was larger than LTS-mediated Ca^2+^ entry in the same location (Egger, 2008; Stroh *et al*., 2012).

If we assume that LTS-mediated Ca^2+^ entry is sufficient to trigger lateral GABA release from at least some reciprocal spines, then the threshold for the generation of dendritic LTS is equivalent to the onset of lateral inhibition, whereas Na^+^-spikes are likely to cause lateral inhibition with greater efficiency than LTS. Pressler and Strowbridge (2017) have predicted that at least 20 coactive MC/TC inputs (within a time window of 1 ms) are required to achieve Na^+^ AP generation in 50% of trials. Because of the rather hyperpolarized GC resting membrane potential of – 80 mV and median unitary EPSP amplitudes ≤ 2 mV (Egger *et al*., 2005; Bywalez *et al*., 2015), we expected a similar outcome.

Another question is whether the local spine spikes contribute to dendritic integration in GCs. Is it conceivable that activation of neighboring spines causes an invasion of the dendritic segment associated with this spine cluster by the spine spike(s)? Could this scenario result in a D-spike? Conventional sequential uncaging (that involves moving the 2D xy-scanner from one uncaging spot to the next) would preclude any observations of such effects because of the inactivation of Na_v_s during the stimulation sequence. Therefore, we implemented a holographic stimulation system to simultaneously stimulate spines in 3D (Go *et al*., 2019). This paradigm is also coherent with physiological activation, since the firing of MC/TCs within a glomerular ensemble is precisely locked to the sniff phase and thus can be synchronized within 1 ms (Shusterman *et al*., 2011). Holographic stimulation also enabled us to target sufficient numbers of inputs, a problem in 2D because of the low GC spine density (1-2 spines per 10 µm, Saghatelyan *et al*., 2005).

## Results

To study synaptic integration within GC apical dendrites we mimicked simultaneous MC/TC inputs to a defined number and arrangement of GC spines in the external plexiform layer by 2P uncaging of DNI-caged glutamate (Palfi *et al*., 2018; Bywalez *et al*., 2015) at multiple sites in 3D using a holographic projector (Go *et al*., 2019). GCs in juvenile rat acute brain slices were patch-clamped and filled with Ca^2+^-sensitive dye OGB-1 (100 µM) to record somatic V_m_ and Ca^2+^ influx into a stimulated spine and several dendritic locations by 2P Ca^2+^ imaging within a 2D plane (see Methods).

### Subthreshold dendritic integration

To characterize subthreshold dendritic integration in terms of somatic V_m_ we first consecutively stimulated single GC spines to obtain single synapse uEPSPs, followed by simultaneous activation of the same spines, resulting in compound cuEPSP. The number of coactivated spines was increased until either AP threshold or the available maximum were reached (10-12 spines, see Methods). Under the given experimental conditions, we succeeded to elicit an AP in 35 out of 111 GCs, 8 of which fired 2 or more APs (e.g. Fig 1a), with an average latency of 41 ± 40 ms (average ± SD, also below). The average single spine uEPSP across all spiking GCs was 1.4 ± 0.8 mV. Integration was quantified by plotting the arithmetic sum of the respective single spine uEPSP amplitudes versus the actually measured multi-spine cuEPSP amplitude for increasing numbers of coactivated spines, yielding a sI/O relationship for each GC.

**Fig 1.**
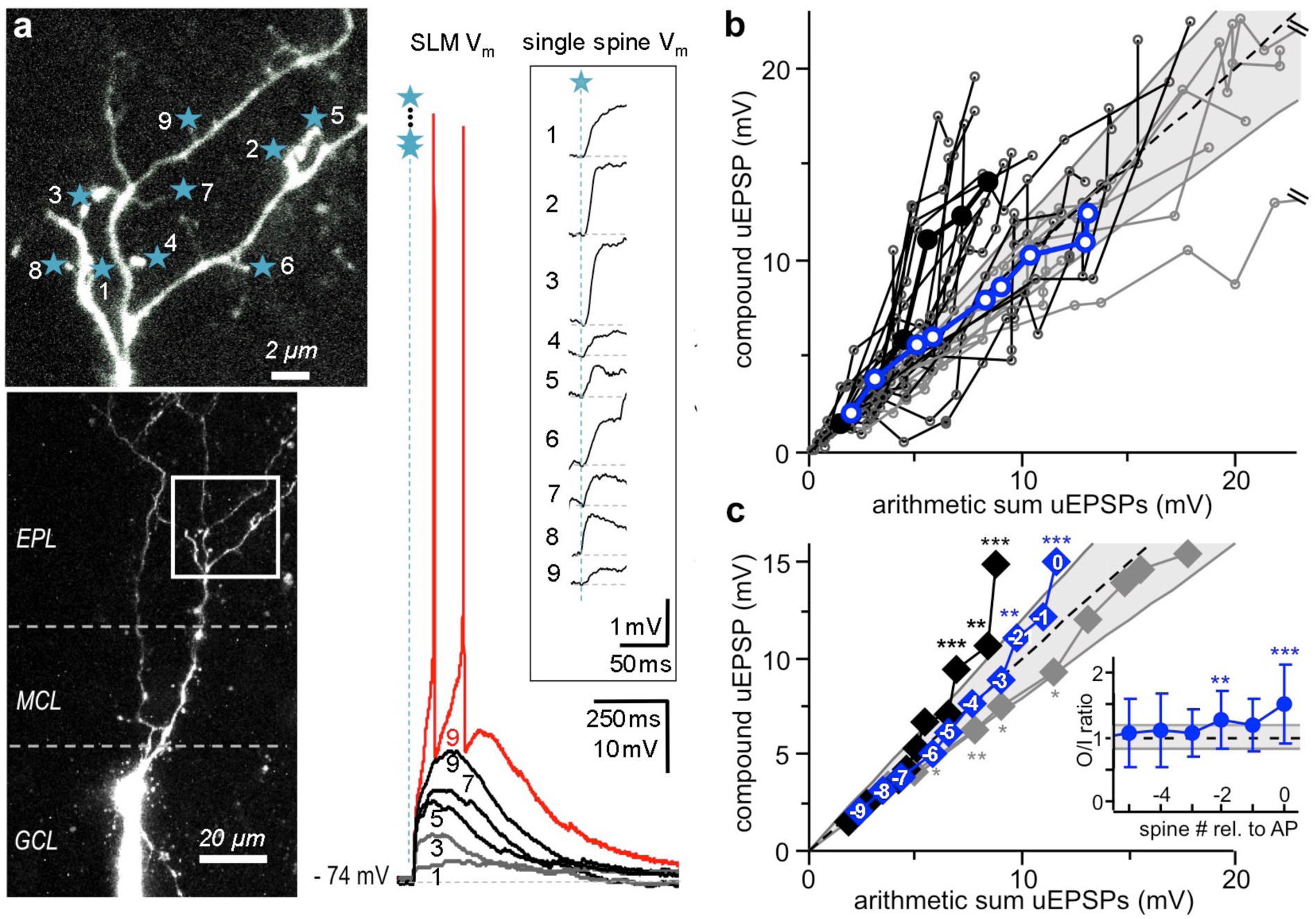
Subthreshold dendritic integration in GCs. **a:** Left: 2P scan and z-projection of representative GC, inset shows enlarged part. Stars represent uncaging spots. Right: Somatic compound uEPSPs and APs generated by simultaneous activation of 1, 3, 5, 7, 9 spines (● in b). Inset: Single spine uEPSPs recorded at the soma (see Methods). In all figures: GCL: granule cell layer, MCL: mitral cell layer, EPL: external plexiform layer. **b**: Subthreshold input/output relationships (sI/O) of n=29 individual experiments in 28 GCs. Grey lines and ○: Sublinear to linear integration. Black lines and circles ○: Supralinear integration. Blue lines and o: Averaged sI/O of 1 to 9 coactivated spines across all GCs. Dashed line: linear y = x. Grey lines: Cut-off supra- and sublinear regime for classification of cells (y = 1.2x, y = 0.8x, see Methods). 2 data points of 2 experiments exceed the scale. **c:** sI/O cumulative plot of experiments in b with data arranged from -9 to 0 spines relative to global AP threshold. Significance levels refer to O/I ratio distributions with means beyond the linear regime (0.8-1.2) tested against linearity (see inset, Methods). ◆: average sI/O of all experiments, statistically significantly supralinear beyond -3 spines (see inset; average O/I ratios for supra- and linear/sublinear sI/Os not shown for clarity): -2 (p = 0.006) and 0 spines (p < 0.001, mean O/I ratio 1.53 ± 0.63). ◆: average of supralinear sI/Os only (n = 19), statistically significantly different from linear summation beyond -3 spines: -2 spines (p < 0.001), -1 (p = 0.007), 0 (p<0.001, mean O/I ratio 1.86 ± 0.52), n=19. ◆: average of sublinear to linear sI/Os only (n = 10), significantly below linear summation below -3 spines: -7 spines (p=0.027), -6 spines (p = 0.008), -5 spines (p=0.02), -4 spines (p=0.021, mean O/I ratio 0.79 ± 0.37), n = 10. In all figures, * p<0.05, ** p<0.01, *** p<0.001.

Fig 1a shows a representative GC where 9 coactivated spines generated an AP in 4 out of 7 trials. This stochastic behavior at AP threshold was also observed in all other spiking GCs in our sample. The sI/O plots of all spiking GCs (Fig 1b) indicate that:

1. For low numbers of coactivated spines, the average sI/O relationship across GCs was linear.
2. In most GCs the cuEPSP amplitude exceeded the arithmetic sum of the single spine uEPSPs beyond a certain stimulation strength by an O/I ratio of at least 1.2 (n = 19 of 29 GCs). We classified these sI/O patterns as supralinear (for further validation of the supralinearity criterion see below and Methods). In this subset of cells, supralinearity was attained at an average of 6.7 ± 2.6 stimulated spines and maintained beyond this threshold until AP generation (except for 1 cell where the last added uEPSP was very large).
3. Consistent sublinear integration (O/I ratio < 0.8) was observed in only 1 GC, while the remaining 9 GCs did not show any consistent deviations from linear behavior. In this subset of cells, the average single EPSP amplitude was larger than for supralinear cells (2.1 ± 0.6 mV vs 1.1 ± 0.6 mV, p < 0.001).

Since each GC required its individual spine number to reach the threshold for AP generation (for the respective pattern of stimulation), we next aligned the sI/O relations to the onset of the global AP before averaging (Fig 1c; see Methods). The ensuing average sI/O relationship was essentially linear until AP threshold ([0], corresponding to the number of spines that in a subset of stimulations triggered an AP) where it turned supralinear. The O/I ratios averaged across cells became significantly supralinear for the first time already at -2 spines below threshold (see Fig 1c inset; see Methods). If only the sI/Os classified as supralinear (see above) were averaged in this way, the average O/I ratio was highly significantly supralinear from 2 spines below AP threshold upwards. The average of the remaining linear/sublinear sI/Os was essentially linear, with a tendency towards sublinearity for lower numbers of coactive spines. Thus, we find that dendritic V_m_ integration is by and large linear at the GC soma, with a supralinear increase in V_m_ close to global Na^+^-spike threshold in the majority of cells.

### Transition from local spine spikes to non-local signals (Ca^2+^- and Na^+^-spikes)

Since GCs are known to feature global LTS and LTS generation had been associated with an increase in EPSP amplitude and duration (Egger *et al*., 2005), we investigated whether the onset of the supralinearity in somatic V_m_ observed in the majority of GC sI/Os coincided with Ca^2+^-spike generation. We detected the transition from local spine spikes (which do not cause detectable dendritic calcium transients, Egger *et al*., 2005; Bywalez *et al*., 2015) to Ca^2+^-spike generation via 2P Ca^2+^ imaging in dendritic shafts that were on average 4.4 ± 3.3 µm remote from the base of the closest stimulated spine, thus not directly adjacent to the spines (e.g. Fig 2a, 4a). Dendritic Ca^2+^ transients were considered to indicate the presence of a Ca^2+^-spike if their amplitude was well above noise level (ΔF/F ≥ 8%, see Methods). We also always imaged a spine that was photostimulated throughout all spine combinations (termed S1 in the following). Fig 2a shows a representative experiment with somatic V_m_ and concurrent Ca^2+^ transients within the S1 spine and at several dendritic locations with increasing numbers of stimulated spines.

**Fig 2.**
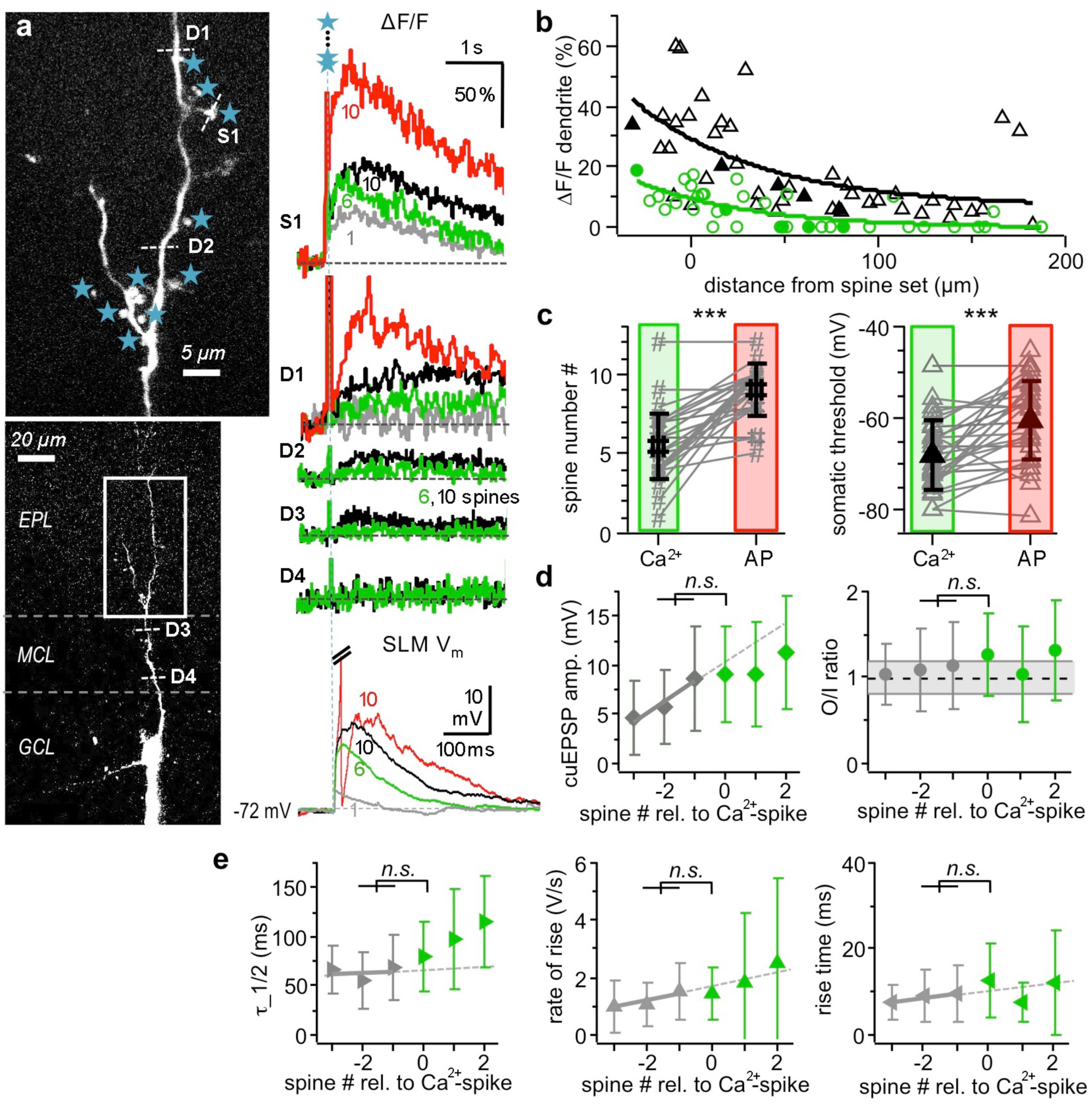
Dendritic Ca^2+^-spikes: A non-local mode of dendritic activation. **a:** Scan of representative GC (left, S1 and D indicate line scan sites, stars indicate uncaging spots, scale bars: 5 µm, 20 µm) with somatic V_m_ traces of single spine and multisite uncaging (bottom right, Na^+^-AP truncated) and averaged (ΔF/F)_TPU_ in the spine and dendrite (top right) upon activation of 1, 6, 10 and 10 spines. Averaged (ΔF/F)_TPU_ in the dendrite measured at increasing distance from the activation site upon subthreshold activation of 6 and 10 spines (middle right). Grey: signals subthreshold for Ca^2+^-spike, green: at Ca^2+^-spike threshold, black: EPSPs and associated ΔF/F signals at AP threshold, red: suprathreshold for global Na^+^-AP, S1: line scan spine S1, D1-4: line scans dendrite D1-4. **b:** Dendritic Ca^2+^ signals versus distance from the center of the stimulated spine set.○: responses at Ca^2+^-spike threshold, Δ: responses for EPSPs at Na^+^-spike threshold. Data from 12 GCs with ΔF/F data imaged at various distances from the set of stimulated spines. Solid symbols: Data from cell in **a**. Green and black lines: Exponential fits to respective data sets (at Ca^2+^-spike threshold: decay constant ± SD: λ = 61 ± 30 µm, ΔF/F(200 µm) = 0 %, n = 38 data points; at Na^+^-spike threshold: λ = 69 ± 47 µm, ΔF/F(200 µm) = 8 %, n = 44 data points). **c:** Comparison of spine numbers (left) and somatic thresholds (right; both n = 27, p < 0.001, paired t-test) for Ca^2+^-spikes and Na^+^-spikes. **d:** Left: Mean somatic cuEPSP amplitudes with spine numbers aligned relative to Ca^2+^-spike threshold (0; n = 25). Difference between -2/-1 and 0 not significantly different from extrapolated linear fit (p = 0.29; Wilcoxon test, see Methods, see Fig S1a for data points from individual experiments). Right: Mean O/I ratios aligned relative to Ca^2+^-spike, not significantly different from subthreshold (p=0.78, n = 25). Grey symbols: subthreshold Ca^2+^-spike, green symbols: suprathreshold Ca^2+^-spike, dashed line: linear fit of subthreshold mean amplitudes, also for **e**. **e:** Kinetics of cuEPSPs (n = 25 GCs, see Fig S1a for data points from individual experiments): No significant increase above extrapolated linear fits at Ca^2+^-spike threshold for half duration (left, p = 0.42, n = 22), rate of rise (middle, p = 0.052, n = 25) or rise time (right, p = 0.49, n= 25).

**Fig 3.**
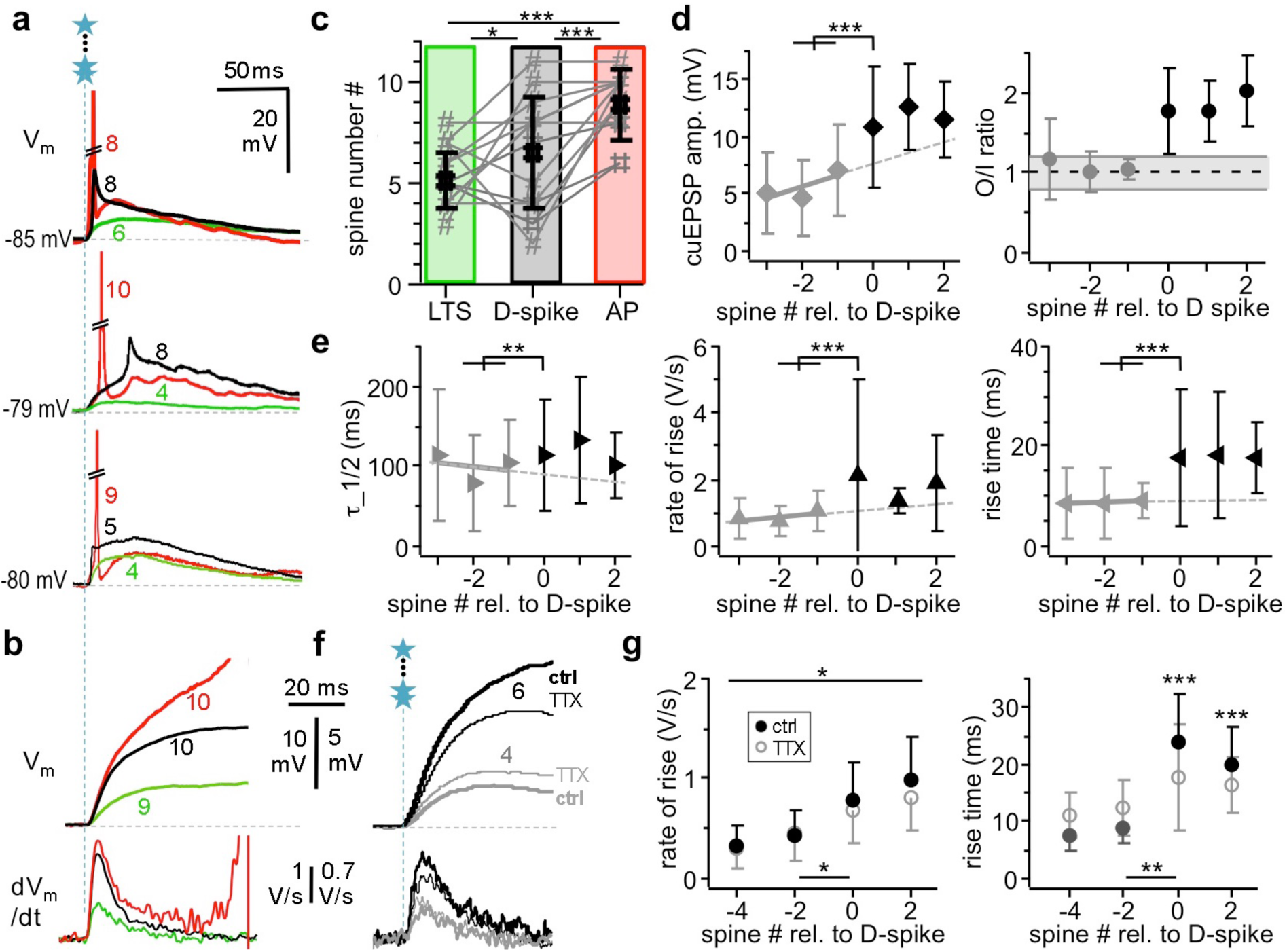
Dendritic Na^+^-spikes. **a:** Examples of somatic spikelets recorded from 3 different GCs at different number of coactivated spines. Green traces and spine numbers: Ca^2+^-spike threshold. Black traces and numbers: Spikelets. Red traces and numbers: Full blown Na^+^ APs, truncated. **b:** Example cuEPSP V_m_ recording in rising phase (top) and 1^st^ derivative (bottom). Colors as in a, black here: D-spike, indicated by increase in rate of rise. **c:** Comparison of spine numbers sufficient to elicit Ca^2+^-spike, D-spike and Na^+^-AP in the same GCs (n = 18 GCs; F_(2,53)_=20.753, p < 0.001, Ca^2+-^spike: 5.1±1.4, D-spike: 6.5 ± 2.7, AP: 8.9 ± 1.7 Holm-Sidak post hoc: Ca^2+-^spike vs D-spike: p = 0.027, Ca^2+-^spike vs AP: p < 0.001, D-spike vs AP: p<0.001). **d:** Left: Mean somatic cuEPSP amplitudes with spine numbers aligned relative to D-spike threshold for GCs with supralinear sI/Os (n = 18). Difference between -2/-1 and 0 highly significantly different from extrapolated linear fit (p = 0.0001; Wilcoxon test, see Methods, see Fig S1b for data points from individual experiments). Right: Mean O/I ratios aligned relative to D-spike. Increase from 1.03 ± 0.13 to 1.77 ± 0.54, not tested, since onset of supralinear O/I ratios was criterion for D-spike threshold. **e:** Kinetics of cuEPSPs (n = 18 GCs; spikelet data included): Highly significant increases beyond extrapolated linear fits at D-spike threshold for half duration (left, p = 0.009, n = 16), rate of rise (middle, p = 0.002, n = 18) or rise time (right, p < 0.001, n = 18). See Fig S1b for data points from individual experiments. **f:** Example for effect of TTX on cuEPSP kinetics below and at D-spike threshold (4 and 6 coactivated spines in this GC, respectively). Top traces: V_m_, bottom: dV_m_/dt, as in **b**. Grey traces: below D-spike threshold, black traces: at D-spike threshold. Thick lines: control, thin lines: in presence of 0.5 – 1 µM TTX. Note the reduction in maximal rate of rise at threshold but not subthreshold. **g:** Cumulative data for effect of TTX (n = 7 GCs with supralinear sI/Os) on cuEPSP rate of rise (left) and rise time (right). Repeated measures two-way ANOVA (see Methods, also below): no interaction effect on rate of rise (spine # x TTX): F_(3,55)_=3.058, p=0.055; TTX effect: F_(1,55)_=8.249, p=0.028. Interaction effect on rise time (spine # x TTX): F_(3,55)_=12.487, p<0.001. Asterisks indicate significance of differences between TTX and ctrl (* p=0.028, *** p<0.001). Asterisks at bottom indicate significance of differences of parameter increases from -2 to 0 between ctrl and TTX (Wilcoxon test; rate of rise: p < 0.05 (W = 17, n_sr_ = 6), rise time: p < 0.01 (W=28, n_sr_ = 7)). See Fig S1c for individual data points.

**Fig 4.**
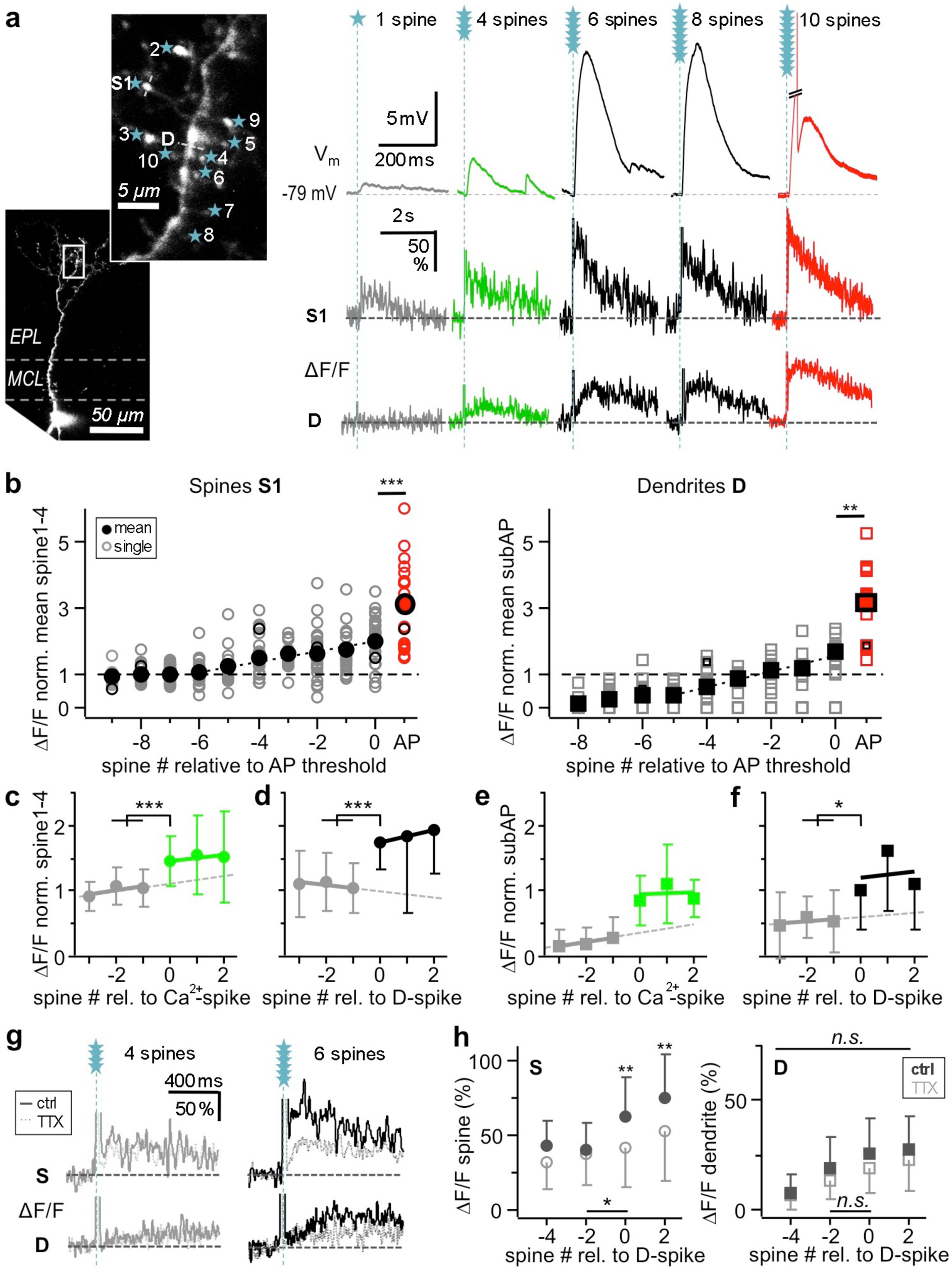
Additional ΔCa^2+^ in spine and dendrite due to non-local spikes. **a:** Scan of representative granule cell (left, S1 and D indicate line scan sites, stars indicate uncaging spots, scale bars: 5 µm, 50 µm) with somatic V_m_ recordings (top right, scale bars: 5 mV, 200 ms) and line scans in spine S1 and dendritic location D for increasing spine numbers (bottom right, scale bars: 50 %, 2 s). Grey traces: below Ca^2+^-spike, green traces: at Ca^2+^-spike threshold; black traces: at and above D-spike threshold; red traces: suprathreshold for global Na^+^-spike. APs cut-off for clarity. **b:** Left: Spine Ca^2+^ signals (ΔF/F)_TPU_ normalized to S1-4 (see Methods) and aligned to AP threshold (n = 33 spines in 16 GCs); ○: individual spines,●: mean; ○: spine from **a**, ○: AP data from individual spines; ●: mean AP data (responses with more than 1 AP were not taken into account). Linear increase from -6 spines onwards (linear fit of mean values until 0: r^2^ = 0.955, p < 0.001) of spine (ΔF/F)_TPU_ with highly significant additional increase upon AP generation (n = 20 pairs, Wilcoxon test, p < 0.001). Right: Dendritic Ca^2+^ signals (ΔF/F)_TPU_ normalized to mean above Ca^2+^-spike threshold (see Methods; n = 19 GCs). Symbols as in right panel, with squares instead of circles. Linear increase from -5 spines onwards (linear fit of mean values: r^2^= 0.879, p<0.001) with significant additional increase upon AP generation (n = 12 pairs, Wilcoxon test, p=0.001). **c:** Spine (ΔF/F)_TPU_ normalized as above, arranged relative to Ca^2+^-spike threshold (n = 26 in 14 GCs, Wilcoxon test, see Methods, p < 0.001) and in **d** relative to D-spike threshold (n = 19 in 10 GCs, P < 0.001). **e:** Dendrite (ΔF/F)_TPU_ normalized as above, arranged relative to Ca^2+^-spike threshold (left: n = 17, significance not tested, since increase in dendritic ΔF/F above noise level was criterion for Ca^2+^-spike) and in **f** relative to D-spike threshold (n = 12, Wilcoxon test, p = 0.015). **g:** Example for effect of 0.5 – 1 µM TTX on ΔF/F below and at D-spike threshold (4 and 6 coactivated spines, respectively; same cell as in Fig 3f). Grey traces: below D-spike threshold, black traces: at D-spike threshold. Solid lines: control, dotted lines: TTX. Note the reduction in spine (ΔF/F) by TTX at threshold but not subthreshold. **h:** Cumulative data for effect of 0.5 -1 µM TTX (n = 7 GCs with D-spike) on (ΔF/F) in spines (left, n = 13) and dendrite (right, n = 7). Full symbols: control, open symbols: TTX. Repeated measures two-way ANOVA (see Methods, also below): interaction effect on spine (ΔF/F) (spine # x TTX): F_(3,103)_=3.195, p=0.035. No interaction effect on dendrite (ΔF/F): F_(3,55)_=0.659, p=0.588; no TTX effect: F_(1,55)_=5.106, p=0.065. Asterisks indicate significance of differences between TTX and ctrl (**: p < 0.01). Asterisks at bottom indicate significance of differences of ΔF/F from -2 to 0 between ctrl and TTX (Wilcoxon test; spine S: p = 0.029 (W = 55, n_sr_ = 13), dendrite: not significant (W=-2, n_sr_ = 5)). See Fig S1c for individual data points.

Across all GCs that could produce both Ca^2+^- and global APs under our experimental conditions, stimulation of on average 5.5 ± 2.1 spines sufficed for Ca^2+^-spike generation (at an average somatic threshold of -67.8 ± 7.6 mV), whereas activation of 9.0 ± 1.6 spines was required to elicit an AP (at a somatic threshold of -60.2 ± 8.8 mV; both spine number and threshold: p < 0.001 Ca^2+^- vs Na^+^-spike, Fig 2c). We also investigated Ca^2+^-spike thresholds in 25 GCs that did not yet fire a Na^+^-spike at the maximum number of stimulated spines, which showed no significant difference to those in spiking GCs (5.3 ± 2.3 spines and -72.1 ± 4.3 mV, respectively). Thus, Ca^2+^-spike generation required substantially lower numbers of coactive excitatory inputs than global AP generation. However, when cuEPSPs were aligned to Ca^2+^-spike threshold spine number before averaging (Fig 2d), there was no discontinuous increase in amplitude at threshold (i.e. not significantly different from linear fit to subthreshold regime, see Methods), and also no significant increase in O/I ratio. Thus, the onset of a Ca^2+^-spike as reported by dendritic ΔF/F is not substantially involved in the generation of V_m_ supralinearity.

The kinetics of the cuEPSP also did not change significantly at Ca^2+^-spike threshold (Fig 2e). Aside from the lack of EPSP boosting and broadening, there was another discrepancy between the Ca^2+^-spike reported here and earlier observations, since the LTS generated by glomerular or external electrical field stimulation (Pinato and Midtgaard, 2005) was an all-or-none event that spread evenly throughout the GC dendrite, whereas here dendritic Ca^2+^-spikes attenuated substantially while propagating from the activated spine set along the dendrite towards the soma (Fig 2b). Thus, the Ca^2+^-spike reported here is mostly a regional signal, which also explains the lack of effects on somatic cuEPSP amplitude and kinetics. Beyond the Ca^2+^-spike threshold, higher numbers of activated spines resulted in larger dendritic ΔF/F signals with increased extent (Fig 2a, b; see also Fig 4a), which can be explained by the recruitment of additional voltage-dependent conductances (see below).

Transition to supralinear behavior due to dendritic Na^+^-spikes (D-spikes) Since Ca^2+^-spike onset did not coincide with the transition to supralinear integration, we investigated the transition between linear and supralinear regimes in more detail in the GCs with supralinear sI/Os (n = 18). This transition happened at an average of 6.5 ± 2.7 spines, significantly higher than the spine number required for Ca^2+^-spike onset and lower than the spine numbers for Na^+^-spikes (Fig 3c). Arrangement of the data relative to the transition spine number (Fig 3d) show a significant discontinuous increase in cuEPSP amplitudes at threshold (i.e. significantly different from linear fit to subthreshold regime, see Methods), and a concomitant increase in mean O/I ratios. The alignment of cuEPSP kinetics to the spine number at the transition in amplitude also revealed highly significant increases of half duration τ_1/2, maximal rate of rise and rise time (Fig 3a, b, e).

An increased rate of rise could indicate the occurrence of a dendritic D-spike (Losonczy and Magee, 2006; Zelles *et al*., 2006). Indeed, in 7 GCs (out of 35 spiking cells) distinct spikelets were detected at the soma, a hallmark of attenuated D-spikes (Fig 3a; Golding and Spruston, 1998; Stuart *et al*., 2007; Llinas and Nicholson, 1971; Smith *et al*., 2013; Epsztein *et al*., 2010). However, how can such D-spikes be consistent with the observed increase in overall rise time? Previously, we had observed that single spine uEPSP rise time increased upon blocking of Na_v_s by wash-in of Tetrodotoxin (TTX; Bywalez *et al*., 2015).

This apparent discrepancy can be explained by a substantial latency of D-spikes. While we could not determine the peak of the D-spike in most GCs, the latency between TPU onset and the peak of spikelets was 21 ± 19 ms (median 10 ms; n = 7 GCs). The delayed occurrence of the D-spike increases the total rise time of the ΔV_m_ signal and also indicates that D-spikes are unlikely to be globalized spine spikes (see Discussion).

Finally, we tested whether the observed changes in EPSP kinetics were indeed due to the activation of dendritic Na_v_s. Application of TTX (0.5 - 1 µM) to a set of 7 GCs with supralinear sI/Os significantly reduced both the increases in cuEPSP rise time and maximal rate of rise at supralinearity threshold (Fig 3f, g). Note that below threshold cuEPSP rise times were slowed in TTX, as expected for spine spike-mediated signals (Bywalez *et al*., 2015).

Moreover, there was a significant increase in ΔCa^2+^ both in activated spines and in nearby dendrites associated with the transition to the D-spike in these experiments (see Fig 4d, f), which was also sensitive to Na_v_ blockade (Fig 4g, h). This observation further proves the presence of a Na_v_-mediated D-spike, since dendritic Na_v_ activation will recruit both low-voltage-activated (LVA) and HVA Ca_v_s and thus can further contribute to Ca^2+^-spikes (Egger *et al*., 2003; Isaacson and Vitten, 2003).

In conclusion, the supralinearity observed in the sI/Os of most GCs is due to the onset of a D-spike.

### Additional Ca^2+^ influxes into the spine mediated by Ca^2+^-, D- and global Na^+^-spike

Apart from the D-spike, can Ca^2+^-spikes and global APs also boost Ca^2+^ influx into spines that are already activated by local inputs? Such summation had been observed previously for both synaptically evoked APs and LTS (Egger *et al*., 2005; Egger, 2008).

Fig 4a shows an exemplary transition from local spine activation to Ca^2+^-spike to D-spike to full blown Na^+^-AP, and in Fig 4b all normalized Ca^2+^ signals are arranged relative to Na^+^-AP threshold in both spine S1 and dendrite. Since for low numbers of coactivated spines (1-4, not aligned to AP threshold, not shown) there was no significant difference in the S1 Ca^2+^ signal, we normalized S1 ΔF/F of each GC to its mean of S1 (1-4) to reduce variance (see Methods).

From 5 coactive spines below AP threshold onwards, S1 and dendritic ΔCa^2+^ increased in a linear fashion up to AP threshold. This observation seems to imply that an individual GC spine could ‘know’ about the number of coactive spines from 5 spines below AP threshold upwards, based on the average amount of extra Ca^2+^ entry. However, arrangement of the data relative to Ca^2+^-spike threshold (as detected in the dendrite, Fig 4e) revealed that below threshold spine (ΔF/F) was by and large constant, whereas at threshold a highly significant increase in ΔCa^2+^ occurred in S1 (by on average ± SD: 1.44 ± 0.80 [0]_Ca2+-spike_ vs [-1/-2] _Ca2+-spike_, n = 25 spines, Fig 4c). Similarly, arrangement of the data relative to the D-spike threshold also revealed a highly significant step-like increase in S1 ΔF/F (by 1.75 ± 0.85 [0]_D-spike_ vs [-1/-2]_D-spike_, n = 18, Fig 4d), and a similar trend in dendritic ΔF/F (by 1.76 ± 0.74 [0]_D-spike_ vs [-1/2]_D-spike_, n = 9, Fig 4f). Finally, global AP generation lead to yet more substantial, highly significant additional Ca^2+^ influx into both the spine (2.03 ± 1.11 [AP] vs [0], absolute 84 ± 59 % ΔF/F, n = 18) and the dendrite (2.03 ± 1.12 [AP] vs [0], absolute 41 ± 20 % ΔF/F, n = 11, both Fig 4b). Compared to the local spine spike, global APs increased spine Ca^2+^ entry by 3.08 ± 1.32 (Fig 4b), thus coincident local inputs and global APs are likely to summate highly supralinearly (see Discussion).

From all these observations, we infer that all three types of non-local signals, Ca^2+^-spike, D-spike and global AP, can mediate substantial additional Ca^2+^ influx into the spine on top of the contribution of the local spine spike, and that the apparently linear mean increase in spine ΔCa^2+^ (Fig 4b) is actually due to overlapping step-like increases due to Ca^2+^-spike and D-spike onsets at different spine numbers in individual GCs. Thus, a GC spine ‘knows’ about the general excitation level of the GC, but cannot resolve individual added coactive spines. Similar step-like increases will occur in dendrites close to the activated spine set and also nearby silent spines (not receiving direct inputs), since those were found previously to respond with similar increases in ΔCa^2+^ to non-local spikes as dendrites (Egger *et al*., 2005; Egger *et al*., 2003; Egger, 2008).

### Molecular mechanisms of integration: Na_v_s

We observed previously (Bywalez *et al*., 2015) that single GC spine activation resulted in a local spine spike, mediated by Na_v_s. While most of the postsynaptic Ca^2+^ entry was mediated by NMDARs, the spine spike contributed additional Ca^2+^ by gating of HVA Ca_v_s. Notably, somatic V_m_ was not reduced in amplitude by Na_v_ blockade, only slowed down in its kinetics, indicative of a strong filtering effect by the spine neck and possibly the GC dendrite (Bywalez *et al*., 2015). A key question of our investigations was whether spine spikes themselves could eventually occur simultaneously across a few clustered spines and thus engender non-local spiking?

For all pharmacological interventions related to dendritic integration mechanisms below the global Na^+^-spike threshold we stimulated 1, 2, 4, 6, 8, 10 spines before and after wash-in of the drug (see Methods). We blocked Na_v_s by wash-in of 0.5 - 1 µM TTX (n = 12 GCs, Fig 5 top). Amplitudes of single and cuEPSPs were unaltered (Fig 5b). However, the significant increase of average O/I ratios from 6 to 8 coactivated spines in control was blocked in the presence of TTX (Fig 5c). 4 of the 12 GCs fired an AP upon stimulation of 10 spines, which was always abolished by wash-in of TTX.

**Fig 5.**
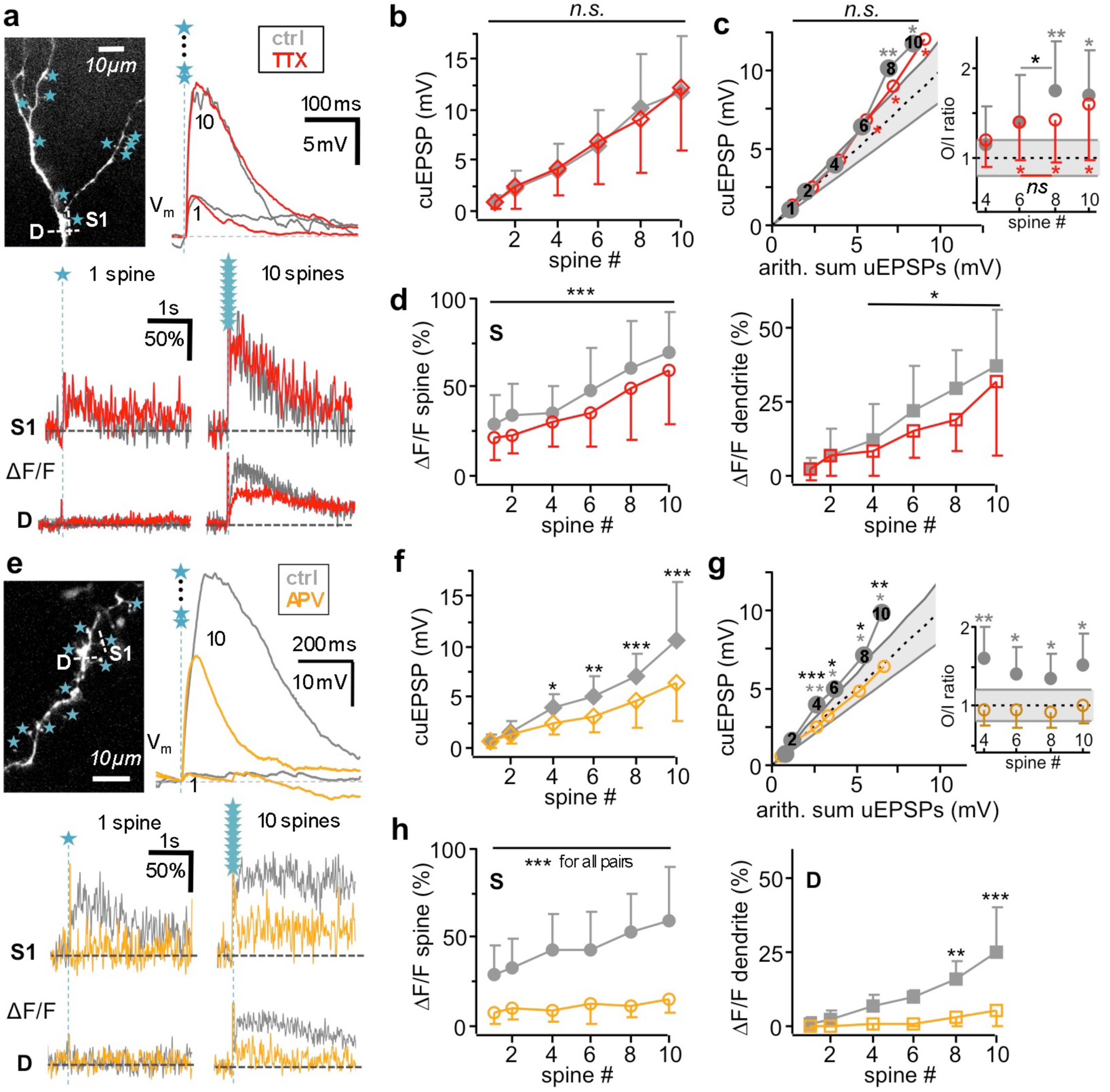
Molecular mechanisms of subthreshold integration: Na_v_s and NMDARs. **a:** Example Na_v_ blockade experiment. Top left: Scan of stimulated spine set with indicated line scan sites S1, D and uncaging spots. Top right: Somatic V_m_ recording of cuEPSP for 1 and 10 coactivated spines, respectively. Bottom: (ΔF/F)_TPU_ in spine (S1) and dendrite (D) for 1 and 10 coactivated spines. Scale bars: 50 %, 1 s. Grey traces: Control. Red traces: TTX (0.5-1 µM). **b:** Average effect (n = 12 GCs) of 0.5-1 µM TTX on somatic cuEPSP amplitude upon activation of 1, 2, 4, 6, 8, 10 spines. Repeated measures two-way ANOVA (see Methods, also below): no interaction effect (spine # x TTX): F_(5,119)_=2.542, p=0.091; no TTX effect: F_(1,119)_=0.965, p=0.352. **c:** Effect (n = 12 GCs) of Na_v_ blockade on averaged sI/O upon activation of 1-10 spines. No interaction effect (spine # x TTX): F(5,99)=1.602, p=0.195; no TTX effect: F_(1,99)_=0.835, p=0.385 (n.s. above data points). Dashed line indicates linear summation, solid grey lines indicate supralinearity (1.2) and sublinearity (0.8). Asterisks/ Significance levels refer to O/I ratio distributions with means beyond the linear regime (0.8-1.2) tested against linearity (as in Fig 1c; see inset, Methods). The average O/I ratio for 8 spines was highly significantly supralinear in control (** above data points, p < 0.01) and significantly increased vs the O/I ratio for 6 spines (* in inset, p < 0.05, Wilcoxon test). While O/I ratios in TTX were significantly supralinear (* below data points, p < 0.05 for all), the increase from 6 to 8 spines disappeared in TTX (inset, n.s.). **d:** Effect of Na_v_ blockade on average spine S (left, n = 25 spines in 11 GCs) and dendrite D (right, n = 12 GCs) (ΔF/F)_TPU_ upon activation of 1-10 spines. No interaction effect on spine (ΔF/F)_TPU_ (spine # x TTX): F_(5,239)_=1.686, p=0.145; TTX effect: F_(1,239)_=15.164, average reduction to 0.89 ± 0.54 of control, p<0.001. No interaction effect on dendrite (ΔF/F)_TPU_ from 4 spines onwards (spine # x TTX): F_(3,79)_=0.457, p=0.715; TTX effect: F_(1,79)_=9.289, average reduction to 0.75 ± 0.28 of control, p=0.014. Asterisks above error bars indicate significance of differences between TTX and control. **e:** Example NMDAR blockade experiment with strong NMDAR signaling component. Top left: Scan of stimulated spine set with indicated line scan sites S1, D and uncaging spots. Scale bar: 10 µm. Top right: Somatic V_m_ recording of cuEPSP for 1 and 10 coactivated spines, respectively. Scale bars: 10 mV, 200 ms. Bottom: (ΔF/F)_TPU_ in spine (S1) and dendrite (D) for 1 and 10 coactivated spines. Scale bars: 50 %, 1 s. Grey traces: Control. Dark yellow traces: APV (25 µM). **f:** Average effect (n = 8 GCs) of 25 µM APV on somatic cuEPSP amplitude upon activation of 1, 2, 4, 6, 8, 10 spines. Interaction effect (spine # x APV): F(5,95)=8.08, p<0.001. Black asterisks indicate significance of differences between APV and control. **g:** Effect of NMDAR blockade on averaged sI/O upon activation of 1-10 spines. Interaction effect (spine# x APV): F_(5,95)_= 3.37, p=0.014, n = 8. Black asterisks above data points indicate significance of differences between APV and control. Dashed line indicates linear summation, solid grey lines indicate supralinearity (1.2) and sublinearity (0.8). Grey asterisks/significance levels refer to O/I ratio distributions with means beyond the linear regime (0.8-1.2) tested against linearity (as in c, see inset, Methods). At control, O/I ratios from 4 spines upwards were supralinear, which all became linear in APV. **h:** Effect of NMDAR blockade on average spine S (left, n = 15 spines) and dendrite D (right, n = 8) (ΔF/F)_TPU_ upon activation of 1-10 spines. Interaction effect on spine (ΔF/F)_TPU_ (spine # x APV): F_(5,179)_=6.36; p<0.001. Interaction effect on dendrite (ΔF/F)_TPU_ (spine # x APV): F_(5,95)_=8.34, p<0.001. Asterisks indicate significance of differences between APV and control. (* p<0.05, **p<0.01, ***p<0.001).

In 7 out of the 12 GCs summation was supralinear, and as shown above (Fig 3), supralinear integration is associated with the occurrence of D-spikes. Indeed, the supralinear increase in V_m_ at threshold was significantly reduced in the presence of TTX (Fig S1c).

Across all 12 GCs the S1 spine Ca^2+^ signal and dendritic Ca^2+^ signals were significantly and similarly reduced in TTX across all numbers of activated spines (Fig 5d; spine: 0.89 ± 0.54 of control, p<0.001; dendrite: 0.75 ± 0.28 of control, p=0.014). Moreover, as shown above, the stepwise increase in spine ΔCa^2+^ at D-spike threshold (Fig 4d) was abolished in TTX (Fig 4g, h). In summary, Na_v_ blockade had only subtle effects on somatic V_m_ summation and thus spine spikes are unlikely to play a major role (see also Discussion). Dendritic Na_v_ activation however underlies the D-spike and the additional Ca^2+^ entry in both spines and dendrites associated with it.

### Molecular mechanisms of non-local spikes: key role of NMDARs

NMDARs have been shown to contribute substantially to local postsynaptic signaling in GCs (Bywalez *et al*., 2015; Isaacson and Strowbridge, 1998; Egger *et al*., 2005) and to also foster the generation of global Ca^2+^-spikes (Egger *et al*., 2005). To investigate the contribution of NMDARs to integration, we blocked NMDARs by wash-in of APV (25 µM) in n = 8 experiments (Fig 5 bottom).

The cuEPSP amplitude was substantially reduced from 4 activated spines onwards (Fig 5e, f). While under control conditions we observed supralinear integration from 4 spines onwards, blocking of NMDARs switched the average sI/O relationship to linear integration (Fig 5g). In 2 experiments cells fired an AP upon stimulation of 10 spines under control conditions and in 1 of these we could record spikelets at the soma upon stimulation of 8 and 10 spines. All were abolished by wash-in of APV.

APV also highly significantly reduced S1 spine Ca^2+^ signals for all stimulation strengths, effectively blocking the on average linear control increase in ΔCa^2+^ (e.g. at 8 costimulated spines spine ΔF/F: 0.25 ± 0.18 of control, p < 0.001; Fig 5h). Moreover, APV strongly reduced dendritic ΔCa^2+^ and thus prevented Ca^2+^-spike generation (Fig 5h; e.g. at 8 spines dendrite ΔF/F: 0.14 ± 0.15 of control, p = 0.003). APV did reduce the half duration of cuEPSPs from 4 spines onwards (interaction effect (spine # x APV): F_(5,95)_=3.202, p=0.017, absolute mean values at 8 coactivated spines: τ_1/2 control 90 ± 53 ms, APV 37 ± 21 ms, data not shown) but did not interfere with fast kinetics, e.g. the maximal rate of rise of the cuEPSP (no interaction effect (spine # x APV): F_(5,95)_=1.616, p=0.182; no APV effect: F_(1,95)_=2.261, p=0.176, n = 8 GCs, data not shown).

Thus, on top of the already known strong contribution of NMDARs to local postsynaptic Ca^2+^ entry, all forms of non-local spikes and their associated Ca^2+^ influxes are highly NMDAR dependent, even though NMDAR activation happens in the electrically isolated spine heads (see Discussion).

### Molecular mechanisms of non-local spikes: contribution of both low and high voltage-activated Ca_v_s to dendritic Ca^2+^ entry

To verify whether distally evoked Ca^2+^-spikes in GC dendrites are mediated by T-type Ca_v_s as observed earlier for global Ca^2+^-spikes evoked by glomerular stimulation (Egger *et al*., 2005), we investigated their contribution to GC multi-spine signals in n = 11 GCs. Wash-in of 10 µM mibefradil did not alter cuEPSPs upon activation of up to 8 spines. Only for 10 spines cuEPSPs were slightly but significantly reduced by on average 0.8 ± 1.4 mV (p=0.01, Fig 6b). Under control conditions activation of 10 spines also lead to supralinear V_m_ summation (Fig 6c), which was reduced by blockade of T-type Ca_v_s. CuEPSP kinetics were unaltered (data not shown). In one experiment an AP was generated upon stimulation of 10 spines under control conditions, which was abolished in the presence of mibefradil.

**Fig 6.**
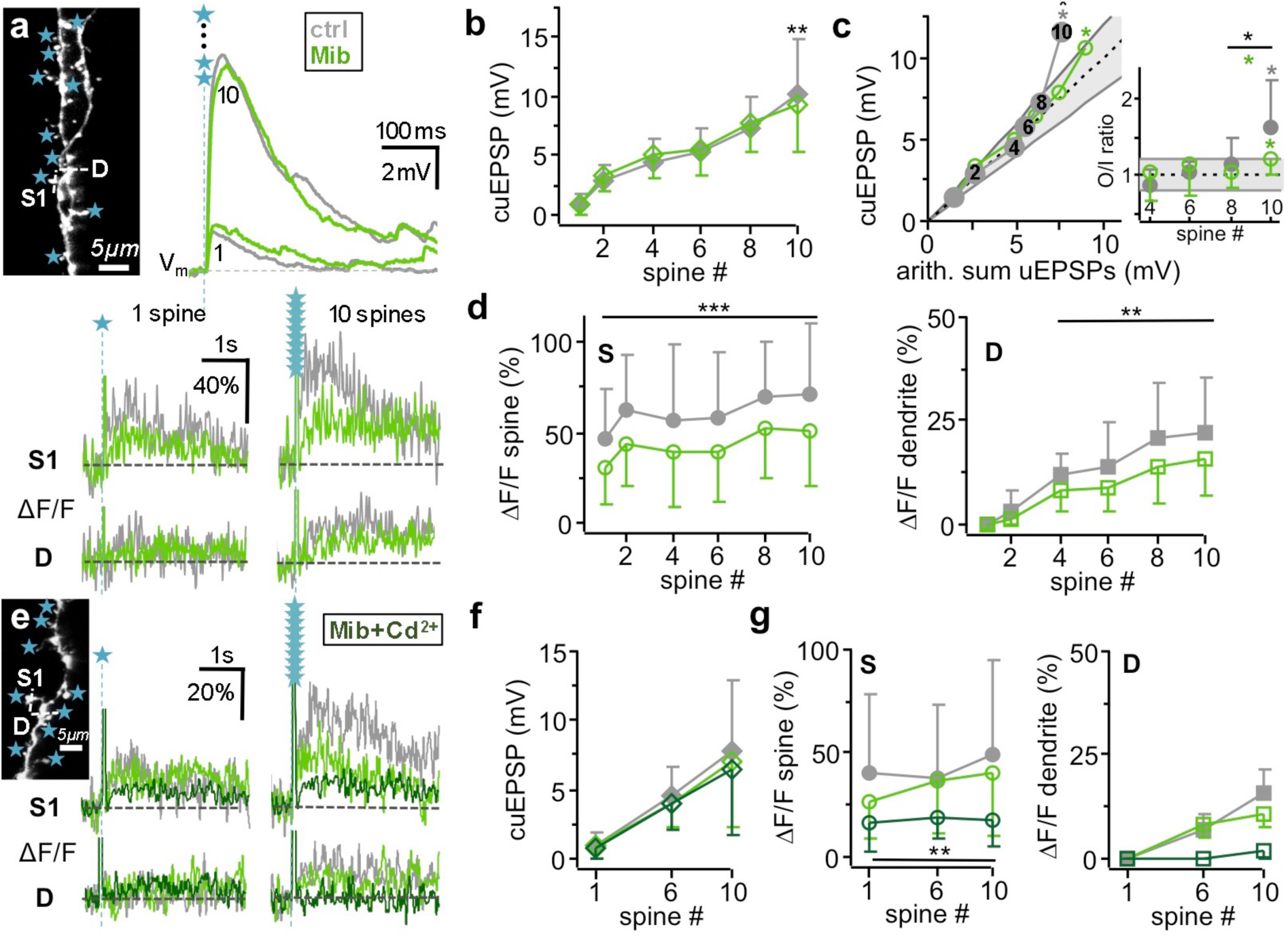
Molecular mechanisms of subthreshold integration: LVA and HVA Ca_v_s. **a:** Example LVA Ca_v_ blockade experiment. Top left: Scan of stimulated spine set with indicated line scan sites S, D and uncaging spots. Top right: Somatic V_m_ recording of cuEPSP for 1 and 10 coactivated spines, respectively. Bottom: (ΔF/F)_TPU_ in spine (S1) and dendrite (D) for 1 and 10 coactivated spines. Grey traces: control. Green traces: Mibefradil (Mib, 10 µM). **b:** Average effect (n = 11 GCs) of 10 µM Mib on somatic cuEPSP amplitude upon activation of 1, 2, 4, 6, 8, 10 spines. Repeated measures two-way ANOVA (see Methods, also below): interaction effect (spine # x Mib) F_(5,131)_=3.88, p = 0.010. Asterisks indicate significance of differences between Mib and control. **c:** Average effect (n = 8 GCs) of LVA Ca_v_ blockade on averaged sI/O upon activation of 1-10 spines. Interaction effect (spine # x Mib) F_(4,69)_=4.695, p=0.006. Black asterisks above error bars indicate significance of differences between Mib and control. Dashed line indicates linear summation, solid grey lines indicate supralinearity (1.2) and sublinearity (0.8). Grey and green asterisks/significance levels above error bars refer to O/I ratio distributions with means beyond the linear regime (0.8-1.2) tested against linearity (*,* p<0.05, as in Fig 1c; see inset, Methods). Integration was significantly supralinear at 10 spines for both control and Mib, as well as the increase in O/I ratios between 8 and 10 spines. However, the O/I ratio increase between 8 and 10 spines in Mib was significantly smaller than for control (inset, Wilcoxon test). **d:** Effect of LVA Ca_v_ blockade on average spine S1 (left, n = 26 spines in 11 GCs) and dendrite D (right, n = 11) (ΔF/F)_TPU_ upon activation of 1-10 spines. No interaction effect on spine (ΔF/F)_TPU_ (spine # x Mib): F_(5,311)_=0.261, p=0.933; Mib effect: F_(1,311)_=60.16, p<0.001. No interaction effect on dendrite (ΔF/F)_TPU_ (spine # x Mib): F_(3,87)_=1.114, p=0.359; Mib effect: F_(1,87)_=15.84, p=0.003. Asterisks indicate significance of differences between Mib and control. **e:** Example for subsequent blockade of LVA and HVA Ca_v_s. (ΔF/F)_TPU_ in spine (S1) and dendrite (D) for 1 and 10 coactivated spines. Top left inset: Scan of stimulated spine set with indicated line scan sites S1, D and uncaging spots. Scale bar: 5 µm. Grey traces: Control. Green traces: Mib (10 µM). Dark green: added Cd^2+^ (100 µM). **f:** Effect of subsequent LVA and HVA Ca_v_ blockade on somatic cuEPSP amplitude upon activation of 1, 6 and 10 coactivated spines (n = 4 GCs). **g:** Effect of subsequent LVA and HVA Ca_v_ blockade upon activation of 1, 6 and 10 spines on spine (ΔF/F)_TPU_ (left, n = 8 spines in 4 GCs) and dendrite (ΔF/F)_TPU_ (right, n = 4). No interaction effect of Cd^2+^ wash-in after Mib on spine (ΔF/F)_TPU_ (spine # x Cd^2+^): F_(2,47)_=1.514, p=0.254; Cd^2+^ effect: F_(1,47)_=14.02, p=0.007. Asterisks indicate significance of differences between Mib+Cd^2+^ and Mib. (* p < 0.05, ** p<0.01, *** p<0.001)

Ca^2+^ signals in spine S1 and dendrite were significantly reduced for all spine numbers (spine: average ± SD 0.74 ± 0.31 of control, p < 0.001; dendrite: 0.74 ± 0.38 of control, p=0.003, Fig 6d). However, mibefradil did not entirely block dendritic ΔCa^2+^ upon stimulation of 4 spines and beyond (remaining signal 16 ± 9 % ΔF/F at 10 coactivated spines).

To identify the source for the remaining dendritic ΔF/F_TPU_, we additionally washed in 100 µM Cd^2+^ to block HVA Ca_v_s in 4 cells (Isaacson and Vitten, 2003). Cd^2+^ effectively abolished the dendritic Ca^2+^ signal and substantially further reduced the S1 spine Ca^2+^ signal to 0.52 ± 0.26 of mibefradil or 0.41 ± 0.22 of control (n = 8 spines), leaving the cuEPSP unaltered (Fig 6 e-g). We conclude that T-type Ca_v_s substantially contribute to Ca^2+^ entry into the spine and dendrite during dendritic integration and mediate the onset of the Ca^2+^-spike, but that HVA Ca_v_s also contribute, most likely involving additional Ca^2+^ entry via L-type Ca_v_s or other channel types that are activated by D-spikes. Both LVA and HVA Ca_v_s did not substantially influence somatic ΔV_m_ in our stimulation paradigm.

### Limited influence of morphology on non-local spike generation

To determine whether the spacing of stimulated spines, the location of the stimulated spine set on the dendrite relative to the MCL, the average spine neck length and other morphological parameters influenced the efficacy of activated subsets of spines to elicit non-local spiking, we analyzed the positions of the stimulated spines relative to the GCs’ dendritic tree as reconstructed in 3D (Fig 7, see Methods). Table S1 shows that only 2 out of 9 morphological parameters correlated with Ca^2+^-spike threshold in terms of coactivated spine numbers, whereas both D-spike and global Na^+^-spike initiation threshold spine numbers did not correlate significantly with any morphological parameter, with a weak trend for a positive correlation between spine distribution and global Na^+^-spike initiation (Fig 7c). Ca^2+^-spike generation was facilitated by close packing of spines that were located on the same and/or a rather low # of branches (Fig 7 c, d).

**Fig 7.**
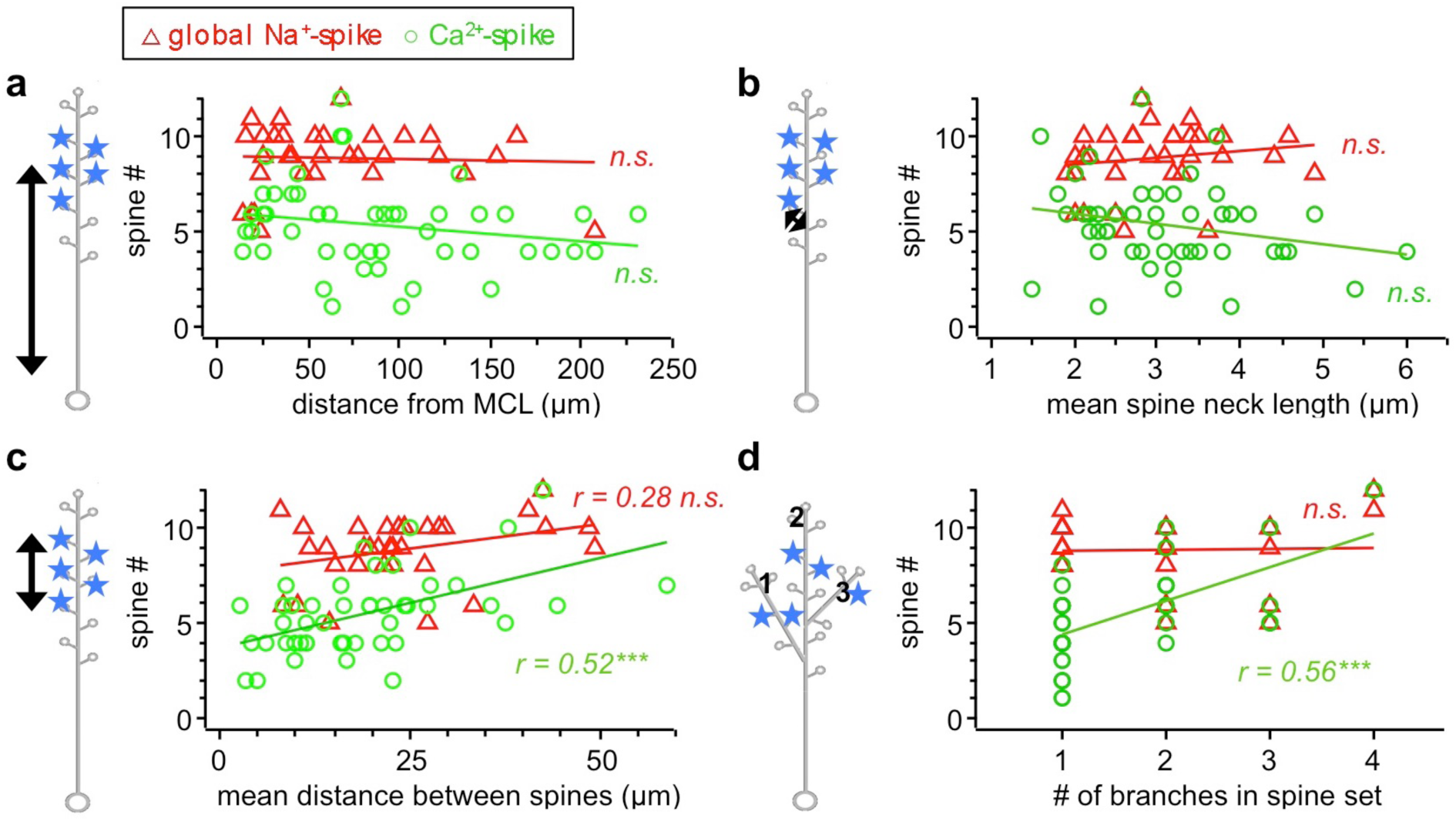
Impact of morphological parameters on threshold spine number for Ca^2+^-spike and Na^+^-spike generation. Ca^2+^-spike data (n = 47 spine sets) are denoted by ○ and global Na^+^-spike data (n = 31 spine sets) by Δ. D-spike data not shown for clarity (but see Table S1). Linear correlation indicated by correlation coefficient r. See Table S1 for power of regressions. **a:** Influence of mean spine distance from the mitral cell layer (MCL) on spine # to elicit Ca^2+^-spikes (r^2^ = 0.019, p = 0.173) and global Na^+^-spikes (r^2^ = 0, p = 0.892). **b:** Influence of the mean spine neck length of activated spine sets on spine # to elicit Ca^2+^-spikes (r^2^ = 0.030, p = 0.120) and global Na^+^-spikes (r^2^ = 0, p = 0.35). **c:** Influence of the spatial distribution of activated spines on spine # to elicit Ca^2+^-spikes (r^2^ = 0.26, p < 0.001, n=47) and global Na^+^-spikes (r^2^ = 0.077, p=0.064). **d:** Influence of number of different dendritic branches that the spine set is distributed across on spine # to elicit Ca^2+^-spikes (r^2^ = 0.31, p < 0.001, n = 47) and global Na^+^-spikes (r^2^ = 0, p = 0.91). (*** p < 0.001).

Within the experimentally accessible range of parameters, individual spine sets have thus by and large an equal impact on local and global Na^+^-spike generation, independent from GC morphology or their relative location on the dendritic tree, which indicates a highly compact GC dendrite and strong isolation of the spines. For Ca^2+^-spike generation clustered spines are more efficient than distributed inputs.

## Discussion

### High excitability of GC apical dendrites

Upon holographic simultaneous multi-spine stimulation, GC dendrites can generate Ca^2+^-, D- and Na^+^-spikes already at rather low numbers of coactivated dendrodendritic inputs (Ca^2+^-spike ∼ 5 inputs, D-spike ∼ 7 inputs, global Na^+^-spike ≥ 9 inputs). Thus, GC dendrites are highly excitable, also in comparison to cortical PCs whose AP threshold required a similar number of coactive spines using the very same holographic system (10 ± 1, n = 7 in 4 PCs, Go *et al*., 2019) although the PCs’ resting potential was depolarized versus GCs by ≥ +10 mV. This high excitability is not due to excessive photostimulation, since the average single EPSP amplitude was smaller in our experiments than in earlier reports on MC/TC-GC synaptic transmission (Bywalez *et al*., 2015; Pressler and Strowbridge, 2017).

Our data also demonstrate that the full set of active dendritic mechanisms known from other neurons (see Introduction) can be triggered solely by dendrodendritic MC/TC inputs to the apical GC dendrite. Because of their location close to the MCL the GCs in our sample (see Methods) belong to superficial GCs (or type III), that preferentially synapse on TC lateral dendrites (Nagayama *et al*., 2014). Superficial GCs are reportedly more excitable than deep GCs (Burton and Urban, 2015), thus our results might not generalize to all GC subtypes, possibly explaining the discrepancy with earlier estimates of excitability (see Introduction; Pressler and Strowbridge, 2017).

The somatic AP threshold (∼ -60 mV) was substantially below Na_v_ activation threshold, indicating a distal AP initiation zone. The threshold spine number reported here is a lower limit, since in ∼2/3 of GCs in our sample full-blown APs at the soma could not yet be elicited at the maximal number of 10 - 12 co-stimulated spines (with the available laser power as bottle neck). Morphological parameters did not influence AP thresholds, indicating that the superficial GC’s dendritic tree is electrotonically compact (see also below).

The low threshold spine number seems to match previous observations that uniglomerular stimulation can already fire GCs (Egger, 2008; Schoppa *et al*., 1998; Geramita *et al*., 2016). On the other hand, ∼20 MC/TCs belong to a glomerular column, with a slight lower share of TCs (Panhuber *et al*., 1985; Liu *et al*., 2016; Royet *et al*., 1998). The release probability at these inputs is ∼0.5 (Egger *et al*., 2005) and thus for uniglomerular activation a given GC is unlikely to be fired solely from intracolumnar dendrodendritic inputs, requiring additional activation most likely originating from MC/TC axonal collaterals (Schoppa, 2006a). However, uniglomerular inputs - if clustered - might suffice to elicit local Ca^2+^ spikes, and MC/TC theta bursts as both observed *in vivo* (Fukunaga *et al*., 2012) could also trigger firing of intracolumnar GCs from the distal apical dendrite.

The high excitability of GCs and the frequent supralinear subthreshold integration indicate a strong role for active conductances in dendritic integration.

### Dendritic spiking: D-spike and localized Ca^2+^-spike

A substantial presence of dendritic Na_v_ channels in GCs was already indicated by a backpropagation study (Egger *et al*., 2003) and recently demonstrated more directly (Nunes and Kuner, 2018). Na^+^ spikelets have not been reported from juvenile rat OB GCs so far; they probably emerged here due to clustered stimulation. In about 2/3 of GCs in our sample we detected D-spikes correlated with the onset of supralinear integration at the soma either as distinct spikelets or, if these were masked by electrotonic filtering, by characteristic step-like increases in the cuEPSP rate of rise, rise time, decay and spine ΔF/F (Fig 3). Significant increases in the EPSP rate of rise and dendritic ΔCa^2+^ indicated D-spikes in CA1 PCs and mouse and frog GCs (Losonczy and Magee, 2006; Burton and Urban, 2015; Zelles *et al*., 2006). The unexpected increase in cuEPSP rise time by almost ∼10 ms can be explained by the substantial latency of spikelets of ∼10-20 ms after TPU onset. This delay indicates that D-spikes are not spatially expanded spine spikes, implying that spine spikes are unlikely to invade the dendrite even under conditions of clustered spine activation, which is further supported by the lack of a correlation between spatial clustering and D-spike or Na^+^-spike threshold spine numbers. Rather, EPSPs are strongly attenuated and also temporally filtered across the spine neck, resulting in slowed integration (Aghvami and Egger, unpublished simulations); moreover, A-type K^+^ currents are known to delay GC firing (Schoppa and Westbrook, 1999) and thus also possibly involved in the yet longer latency of global Na^+^-APs at threshold observed here (∼40 ms). Initiation of D-spikes most likely happens at dendritic Na_v_ hot-spots (Nunes and Kuner, 2018), whereas the existence of a dedicated global Na^+^-AP initiation zone in GC apical dendrites seems probable, with its precise location a matter of speculation at this point (but see Pressler and Strowbridge, 2019).

All GCs in our sample featured Ca^2+^-spikes (in terms of dendritic Ca^2+^ entry), which at threshold were rather regional, decreasing strongly towards the soma. Although Ca_v_ densities are apparently lower in the proximal apical dendrite (Egger *et al*., 2003), Ca^2+^-spikes evoked by glomerular stimulation occurred in an all-or-none fashion throughout the entire dendritic tree with a concomitant increase and broadening of somatic EPSPs that were not observed here(Egger *et al*., 2005). We conclude that the main initiation zone for global Ca^2+^-spikes is probably not located in the distal apical dendritic tree (see also Pressler and Strowbridge, 2019). Thus, in contrast to the global Ca^2+^-spike upon glomerular activation, input to rather densely packed spines might provide a substrate for more local lateral inhibition as suggested earlier (Woolf *et al*., 1991; Woolf and Greer, 1994; Isaacson and Strowbridge, 1998). Anyways, local Ca^2+^-spikes became more global close to Na^+^-AP threshold, along with recruitment of HVA Ca_v_s. Thus, GCs feature multiple levels of compartmentalization.

In contrast to AP generation, Ca^2+^-spike generation is strongly influenced by input distribution, in line with electrotonic attenuation of subtreshhold EPSPs along the dendrite (Tran-Van-Minh *et al*., 2015). Since the Ca^2+^-spike precedes the D-spike and AP and its space constant of ≲ 60 µm covers the maximum spatial extent of spine sets in our experiments, its presence can reduce passive attenuation and thus explain the independence of D-spike and AP generation from input distribution within the spatial regime accessible by our system. Which in turn means that D-spike and global APs are insensitive to spatial changes in local input distributions.

### NMDA-spikes and role of NMDARs in GC synaptic processing

NMDARs contribute substantially to supralinear integration in GCs, both at the level of V_m_ and ΔCa^2+^. They are required for Ca^2+^-spike generation and their blockade had a much stronger effect on V_m_ supralinearity than Na_v_ or Ca_v_ blockade alone. This higher efficiency is probably related to the slower kinetics of the NMDAR component, which therefore is filtered much less both by the spine neck and then along the dendritic tree compared to Na_v_ or Ca_v_ mediated currents (Stuart *et al*., 2016). Conversely, strong filtering of Na_v_ currents explains the small influence of TTX on somatic dendritic integration observed here. The substantial impact of NMDARs on GC dendritic integration is characteristic for NMDA-spikes (Maccaferri and Dingledine, 2002; Antic *et al*., 2010). Accordingly, GC APs evoked by synaptic stimulation are followed by NMDAR-dependent plateau potentials (Egger, 2008; Stroh *et al*., 2012). In most GCs, EPSP half durations were > 50 ms at higher numbers of coactive spines, thus dendritic Ca^2+^- and Na^+^-spikes are closely intertwined with NMDA-spikes.

As a note of caution, holographic uncaging might overemphasize the role of NMDARs, since (1) APV blocks TPU-evoked spine ΔF/F slightly more than synaptic ΔF/F (to 65% vs 50% of control, Bywalez *et al*., 2015) and (2) the axial point spread function of our multi-site uncaging system is extended to 2.7 µm (Go *et al*., 2019) from 1.1 µm, which might activate yet more extrasynaptic NMDARs. However, the effect on APV on single spine (ΔF/F)_TPU_ was similar as in Bywalez *et al*. (2015).

NMDARs are predicted to enable supralinear summation of ΔCa^2+^ at positive Hebbian pairing intervals of single spine spike and global APs (Aghvami *et al*., 2019). The median latency of global Na^+^ APs at threshold of 35 ms observed here matches with the simulated regime of maximally supralinear summation efficiency, which explains the strong increase of ΔCa^2+^ in spines upon AP generation. In conclusion, NMDARs are essentially involved in all aspects of GC reciprocal synaptic processing, including release of GABA from reciprocal spines (Lage-Rupprecht *et al*., 2019) and synaptic plasticity (Chatterjee *et al*., 2016; Neant-Fery *et al*., 2012).

### Functional implications

Our findings imply that dendritic spikes and therewith possibly lateral inhibition can be invoked already at very low numbers of coactive spines. The observed stepwise increases in spine and dendrite ΔCa^2+^ at the three spike thresholds imply that release probabilities for GABA (which is ∼0.3 for local stimulation, Lage-Rupprecht *et al*., 2019) might also be increased in a step-like fashion, thus rendering both lateral and recurrent inhibition more effective. GC-mediated lateral inhibition is thought to implement contrast enhancement and synchronization of gamma oscillations across glomerular columns responding to the same odorant (Urban and Arevian, 2009; Fukunaga *et al*., 2014; Peace *et al*., 2017). Fast gamma oscillations in the bulb are generated at the GC-MC/TC synapse independently of global APs (Lagier *et al*., 2004; Schoppa, 2006b) and require a fast excitatory-inhibitory feedback loop (Pouille *et al*., 2017; Fukunaga *et al*., 2014), that is likely to involve both reciprocal and lateral processing. D-spikes could be powering fast oscillatory lateral and recurrent output as they have shorter latencies than global back-propagating APs. Zelles *et al*. (2006) already proposed interaction of D-spikes and back-propagating APs in GCs at intervals as short as 5-6 ms and also Pinato and Midtgaard (2005) could elicit spikelets at a frequency of 150-250 Hz, whereas the maximum frequency of global GC APs is much lower (10-30 Hz). *In vivo,* GCs display only sparse, long latency AP firing (Cang and Isaacson, 2003; Kato et al., 2012), whereas spikelets have been frequently observed (Wellis and Scott, 1990; Labarrera *et al*., 2013; Mori and Takagi, 1978; Luo and Katz, 2001). Similarly, D-spikes are associated with sharp wave-associated ripples (120-200 Hz) in hippocampal CA1 PCs (Klausberger *et al*., 2004; Kamondi *et al*., 1998; Memmesheimer, 2010).

The supralinear integration in GC dendrites is at variance with that in subtypes of retinal amacrine cells who also release GABA from their dendrites. In these cells dendritic Na_v_s do not amplify EPSPs and compartmentalization is high (Grimes *et al*., 2010), most likely because spatial organization of lateral inhibition in the retina is continuous.Changes in GC spine and dendrite ΔCa^2+^ were not necessarily correlated with changes in somatic V_m_ amplitude (both for the localized Ca^2+^ spike and the attenuated D-spike), allowing for multiplexed signals as proposed for cerebellar GCs (Tran-Van-Minh *et al*., 2016), which in bulbar GCs might implement e.g. independent plasticity induction across reciprocal spines (Chatterjee *et al*., 2016). On a yet more speculative note, different GC spike types might encode for different types of odor information. Such multiplexing of odor information was already described for MCs in zebrafish, where wave-like, most likely GC-mediated, gamma oscillations and tightly phase-locked spiking are tied to odor category and odor identity, respectively (Friedrich *et al*., 2004).

## Material and methods

### Animal handling, slice preparation and electrophysiology

All experimental procedures were done in accordance with the rules laid down by the EC Council Directive (86/89/ECC) and German animal welfare legislation. Rats (postnatal day 11-21, Wistar of either sex) were deeply anaesthetized with isoflurane and decapitated. Horizontal OB brain slices (thickness 300 µm) were prepared and incubated at 33°C for 30 min in ACSF bubbled with carbogen and containing (in mM): 125 NaCl, 26 NaHCO_3_, 1.25 NaH_2_PO_4_, 20 glucose, 2.5 KCl, 1 MgCl_2_, and 2 CaCl_2_. Recordings were performed at room temperature (22 °C). Patch pipettes (pipette resistance 5-7MΩ) were filled with an intracellular solution containing (in mM): 130 K-methylsulfate, 10 HEPES, 4 MgCl_2_, 2.5 Na_2_ATP, 0.4 NaGTP, 10 Na-phosphocreatine, 2 ascorbate, 0.1 OGB-1 (Ca^2+^ indicator, Invitrogen), 0.04-0.06 Alexa Fluor 594 (Life Technologies), at pH 7.3. The following pharmacological agents were bath applied in some experiments: TTX (0.5 – 1 µM, Alomone), D-APV (25 µM, Tocris), mibefradil (10 µM, Tocris), and cadmium chloride (Cd^2+^, 100 µM, Sigma). Drugs were washed in for at least 10 min before starting measurements. Electrophysiological recordings were made with an EPC-10 amplifier and Patchmaster v2.60 software (both HEKA Elektronik). GCs were patched in whole-cell current clamp mode and held near their resting potential of close to –75 mV (Egger *et al*., 2003). If GCs required >25 pA of holding current, they were rejected. In order to provide optimal optical access to the GC apical dendritic tree, patched GCs were located close to the mitral cell layer (MCL).

### Combined two-photon imaging and multi-site uncaging in 3D

Imaging and uncaging were performed on a Femto-2D-uncage microscope (Femtonics). The microscope was equipped with a 60× water-immersion objective used for patching (NA 1.0 W, NIR Apo, Nikon) and a 20× water-immersion objective used for two-photon (2P) imaging and uncaging (NA 1.0, WPlan-Apo, Zeiss). Green fluorescence was collected in epifluorescence mode. The microscope was controlled by MES v4.5.613 software (Femtonics). Two tunable, verdi-pumped Ti:Sa lasers (Chameleon Ultra I and II, respectively, Coherent) were used in parallel, set to 835 nm for excitation of OGB-1 and to 750 nm for uncaging of DNI-caged glutamate (DNI, Femtonics; Chiovini *et al*., 2014). DNI was used in 0.6 mM concentration in a closed perfusion circuit with a total volume of 12 ml and was washed in for at least 10 min before starting measurements. To visualize the spines and for Ca^2+^ imaging we waited at least 20 min for the dyes to diffuse into the dendrite before starting measurements.

Imaging and uncaging beam were decoupled before the entrance of the galvanometer-based 2D scanning microscope using a polarizing beam splitter to relay the uncaging beam to a spatial light modulator (SLM X10468-03, Hamamatsu). Using our custom-written Matlab based SLM software, we next positioned multiple uncaging spots/foci in 3D at a distance of 0.5 *μ*m from the spine heads. Our system allowed for a maximum number of 12 spots in a volume of 70x70x70 µm^3^, usually spines no deeper than ∼ 30 µm were imaged because otherwise uncaging laser power was too much attenuated. The positioning was checked before each measurement and, if necessary, readjusted to account for drift. The holographic projector module is described in detail in Go *et al*. (2019). The uncaging pulse duration was 1–2 ms and the laser pulse power was adjusted individually for each experiment to elicit physiological responses, depending on the depth of the spines (Bywalez *et al*., 2015). For simultaneous multi-site photostimulation, the total uncaging power and the number of foci/spots were kept constant. ‘Superfluous’ foci, i.e. foci that were not needed as stimulation spots at a given time of an experiment, were excluded by positioning them just outside the holographic field-of-view, such that they would fall off the optics and not be projected onto the sample (Go *et al*., 2019). Imaging of uncaging-evoked Ca^2+^ signals in selected spines and dendritic positions within one 2D plane (see below) was carried out as described earlier (Bywalez *et al*., 2015). During simultaneous Ca^2+^ imaging and photostimulation, imaging was started 700 ms before the uncaging stimulus. During uncaging the scanning mirrors were fixed.

In each experiment, single spines were consecutively activated and somatic uncaging EPSPs (uEPSPs) were recorded for each spine separately. Next, successively increasing # of these spines were simultaneously activated and somatic compound uEPSPs (cuEPSPs) were recorded until the GC fired an AP or, in the experiments with focus on subthreshold integration, until a maximum # of 10 activated spines was reached. A subset of spines and dendritic locations located within the same focal plane were chosen for 2P line-scanning to gather Ca^2+^ imaging data. At least one spine, termed S1 in the following, was always located in this imaging plane to gather complete data from activation of a single spine to activation of n spines. Due to the spine density being higher in distal regions and Ca^2+^ imaging being restricted to one focal plane, most dendritic measurements in a distance from the center of the stimulated spine set (Fig 1b) were proximal to the stimulation site. The sequence of the successively more activated spines with respect to their position on the dendritic tree was randomly chosen. However, the low spine density (see Introduction) and the restriction to a volume of 70x70x70 µm^3^ mostly determined the choice of activated spines. Both single spine stimulations and the different combinations of multi-site uncaging were, if possible, performed at least twice and recordings were averaged for analysis.

Since such experiments were performed with up to 40 different stimulation conditions, we decided to increase the spine # by increments of two for some experiments (in particular for pharmacology) in order to limit the experiment duration and thus to ensure a good recording quality.

### Data analysis

Changes in Ca^2+^ indicator fluorescence were measured relative to the resting fluorescence F_0_ in terms of ΔF/F as described previously (Egger *et al*., 2005). Electrophysiological and Ca^2+^ imaging data were analyzed using custom macros written in IGOR Pro (Wavemetrics). Traces contaminated by spontaneous activity were discarded. As described above, multiple (2 or more) recordings of the same stimulation type were averaged and smoothed (box smoothing) for analysis. uEPSP and (ΔF/F)_TPU_ rise times were analyzed in terms of the interval between 20% and 80% of total uEPSP/(ΔF/F)_TPU_ amplitude; uEPSPs and (ΔF/F)_TPU_ half times of decay (τ_1/2) were analyzed in terms of the interval between the peak and 50% of the total EPSP or (ΔF/F)_TPU_ amplitude. The uEPSP maximum rate of rise was determined by the peak of the first derivative of the uEPSP rising phase. The AP threshold was detected via the zero point of the 2^nd^ derivative of the AP rising phase.

Integration was quantified by plotting the arithmetic sum of the respective single spine uEPSP amplitudes versus the actually measured multi-spine cuEPSP amplitude for increasing numbers of coactivated spines, yielding a subthreshold input-output relationship (sI/O; Tran-Van-Minh *et al*., 2015). If the cuEPSP amplitude consistently exceeded the arithmetic sum of the single spine uEPSPs beyond a certain stimulation strength by at least a factor of 1.2, we classified these sI/O patterns as supralinear. If the factor fell consistently below 0.8, we classified these sI/O patterns as sublinear, and the patterns falling between these categories were considered to be linear. The factors were set at 0.8 and 1.2 to exceed potential undersampling errors in cuEPSP amplitudes (see below) and were empirically confirmed by concurrent characteristic changes in cuEPSP kinetics (see Fig 3e).

As criterion for the presence of a Ca^2+^-spike dendritic Ca^2+^ transient amplitudes (ΔF/F)_TPU_ had to exceed a value well above noise level (≥ 8% ΔF/F or factor 1.5 above noise level of 5% ΔF/F) and be detectable at every dendritic line scan located within the section of the dendrite carrying stimulated spines (Fig 2, Fig 4e; Egger *et al*., 2005).

## Data sampling, normalization and alignment

Since for any particular number of coactivated spines we could not perform more than 2 -3 stimulations in the interest of finishing experiments within the average lifetime of GC recordings, the individual cuEPSP measurements might differ from the mean for that particular spine number due to undersampling. For single uEPSPs, a previous data set allows to estimate the variance at the average EPSP amplitude of 1.40 mV in the experiments in this study as 0.39 mV or ∼ 30% (n = 18 spines, linear regression on data with at least 6 uEPSP measurements per spine on the same system from Bywalez *et al*., 2015). On the other hand, stimulations of larger numbers of spines n can be expected to reduce this sampling problem by a factor of √n, since the variabilities of the single spine responses should cancel out, similar to the effect of averaging across repeated stimulations of the same spine. Therefore, we expect a fairly consistent reporting of cuEPSP amplitudes above 5 costimulated spines, even with a low number of stimulations. Still, we set the criterion for supra-/sublinearity in membrane potential (V_m_) rather high (± 20 %), to prevent any erroneous classifications, also since the undersampling errors further propagate to the calculation of output-input (O/I) ratios.

Similarly, the variance in single spine (ΔF/F)_TPU_ is on the order of 6% ΔF/F or ∼ 20% (again derived from Bywalez *et al*., 2015). To compare S1 spine ΔF/F amplitude data across experiments relative to solely local activation of S1, we intended to normalize these to the spine ΔF/F amplitude for local, unitary activation. Because of the undersampling problem, we tested for up to which spine number there was no significant increase in ΔF/F, which was 4 spines (Friedman repeated measures ANOVA on ranks: Χ^2^_F_(3)=4.802, p=0.187). Therefore, we averaged S1 (ΔF/F) for (co)stimulations of 1, 2, 3, 4 spines and used the mean as basal unitary ΔF/F for normalization. Thus, undersampling could be compensated for by this means.

Since dendritic (ΔF/F)_TPU_ was usually detectable only in stimulations above 4 spines in most cases, (ΔF/F)_TPU_ in the dendrite was normalized to the mean of all stimulations below global Na^+^ AP threshold, or the average size of the dendritic Ca^2+^-spike mediated Ca^2+^-signal.

Since each GC required its individual spine number to reach the threshold for the non-local events Ca^2+^-spike, D-spike and global Na^+^-spike (for the respective pattern of stimulation), we aligned the data in relation to the onset of the non-local event (e.g. Fig 2d, e relative to Ca^2+^-spike). Such realignments allow to reveal effects across the sampled cells that otherwise would be smeared out because of cell-specific thresholds, such as recruitment of active conductances near thresholds (Losonczy and Magee, 2006).

### Morphological analysis

GC apical dendrites were reconstructed from 2P fluorescence z-stacks gathered at the end of each experiment, using Neurolucida (MBF Bioscience). Distances were measured along the dendrite. Mean distances of a spine set were analyzed in terms of the average distance of all stimulated spines from e.g. the MCL. The distribution of a stimulated spine set across the dendrite was analyzed in terms of the mean distance of each spine from all other stimulated spines along the dendrite. Spine neck lengths were estimated as described before (Bywalez *et al*., 2015).

### Statistics

Statistical tests were performed in Sigmaplot 13.0 (Systat Software, Inc) or on vassarstats.net. To assess statistical significance levels across spine numbers or threshold V_m_ values for Ca^2+^-spike versus global Na^+^-spike (Fig 2c), data sets were compared using paired t-tests for dependent data sets. Not normally distributed data sets (Shapiro-Wilk Normality Test) were compared using Wilcoxon signed rank tests. To assess statistically significant differences from linear summation in sI/O relation data sets, the distribution of ratios of the measured uEPSP amplitudes/arithmetic sums (O/I ratio) was tested against a hypothesized population mean/median of 1.0 (corresponding to linear summation), using one-sample t-tests or one-sample signed rank tests for not normally distributed data. To assess variation in repeated measure data sets (Fig 3c) repeated measures ANOVA together with all pairwise multiple comparison procedure (Holm-Sidak-method) was performed. For pharmacology experiments (e.g. Fig 5) repeated measures two-way ANOVA together with all pairwise multiple comparison procedure (Holm-Sidak-method) was performed. For statistical analysis of dendritic (ΔF/F)_TPU_ before and after pharmacological treatment just stimulations of ≥ 4 spines were taken into account, since for lower numbers of spines usually no signal was detectable under control conditions.

Due to the increase of spine numbers by increments of 2 in some experiments, averaged data points for a given spine number do not contain the same n of individual measurements across different spine numbers. Even more so when the data were aligned relative to individual spike thresholds (e.g. alignment relative to Ca^2+^-spike threshold in Fig 2d, e), since not all experiments contained data points for the more remote spine numbers +2 or -3. In addition, in experiments with data gaps just before a global spike threshold at spine number x, it is not possible to know whether the spike threshold could have already been reached at x-1 spines (e.g. alignment relative to Ca^2+^-spike in Fig 2 d, e). We accounted for this uncertainty by averaging the data in the continuous experiments for -2 and -1 and used these averaged data for paired comparison of parameters below and at threshold (non-parametric Wilcoxon test). Fig S1 shows the individual data points for all these comparisons normalized to -2/-1. If there was a significant linear increase or decrease with spine number in the parameter in the subthreshold regime (grey dashed lines in Fig 2d, e; 3d, e; 4c-f), the expected increment based on this change was subtracted from the parameter values at threshold before statistical testing for a difference.

To assess statistical significance for linear increase and decrease (Table S1) we performed a linear regression analysis. Given r^2^ values are adjusted r^2^ values.

