## Supplemental Material for "Dendritic integration in olfactory bulb granule cells: Thresholds for lateral inhibition and role of active conductances upon simultaneous activation"

### Supporting information

Table S1. **Regression between coactivated threshold spine numbers for Ca^2+^-spike, D-spike, and global Na^+^-spike and various morphological parameters and input patterns** (see Methods). n: number of analyzed spine sets, r^2^: adjusted coefficient of determination, p-value: two-tailed significance level of regression, COF: coefficient constant. Statistically significant values are highlighted in yellow.

|  | Influence on generation of | | | | | | | | | | | |
| --- | --- | --- | --- | --- | --- | --- | --- | --- | --- | --- | --- | --- |
|  | **Ca^2+^-spike (n = 47)** | | | | **D-spike (n = 20)** | | | | **global Na^+^-spike (n = 31)** | | | |
| **Parameter** | r^2^ | p | COF | power | r^2^ | p | COF | power | r^2^ | p | COF | power |
| **Spine distribution** | 0.257 | <0.001 | 0.092 | 0.970 | 0.000 | 0.601 | 0.041 | 0.074 | 0.077 | 0.064 | 0.051 | 0.457 |
| **Distance from MCL** | 0.019 | 0.173 | -0.008 | 0.274 | 0.000 | 0.952 | -0.001 | 0.029 | 0.000 | 0.844 | -0.001 | 0.039 |
| **Distance from soma** | 0.022 | 0.156 | -0.007 | 0.293 | 0.000 | 0.806 | 0.003 | 0.043 | 0.000 | 0.763 | -0.002 | 0.048 |
| **# of different branches** | 0.310 | <0.001 | 1.713 | 0.993 | 0.045 | 0.186 | -0.846 | 0.260 | 0.000 | 0.914 | 0.035 | 0.032 |
| **# of preceding bifurcations** | 0.000 | 0.988 | 0.004 | 0.026 | 0.000 | 0.329 | -0.514 | 0.160 | 0.000 | 0.366 | 0.259 | 0.144 |
| **spine neck length** | 0.030 | 0.120 | -0.510 | 0.343 | 0.148 | 0.053 | 1.521 | 0.493 | #: 0.000 | 0.355 | 0.361 | 0.149 |
| **Diameter of proximal dendrite** | #: 0.000 | 0.895 | -0.058 | 0.034 | #: 0.017 | 0.266 | 0.864 | 0.196 | #: 0.037 | 0.140 | 0.597 | 0.313 |
| **Distance 1^st^ branchpoint from MCL** | #: 0.000 | 0.923 | 0.001 | 0.031 | #: 0.000 | 0.565 | -0.010 | 0.080 | #: 0.073 | 0.070 | -0.013 | 0.443 |
| **Distance 1^st^ branchpoint from soma** | #: 0.000 | 0.462 | -0.005 | 0.110 | #: 0.000 | 0.828 | 0.003 | 0.040 | #: 0.046 | 0.122 | -0.009 | 0.339 |
| **Single spine uEPSP amplitude** | #: 0.000 | 0.718 | 0.172 | 0.055 | #: 0.074 | 0.130 | -1.683 | 0.326 | #: 0.000 | 0.406 | -0.333 | 0.128 |

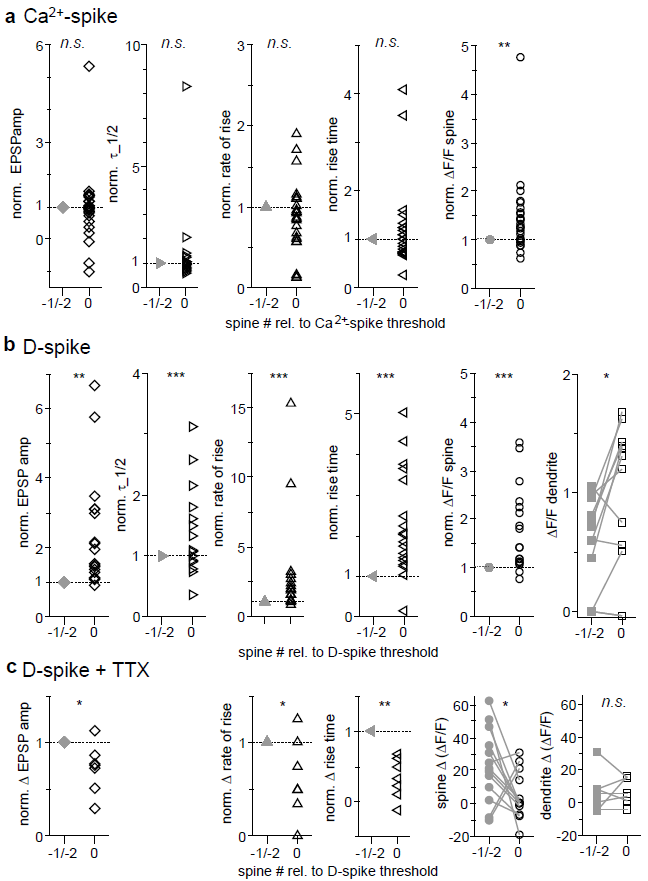

Fig S1. **Individual data sets at threshold for Ca^2+^-spike and D-spike**

Individual data points from paired data comparisons across threshold for Ca^2+^-spikes (**a**), D-spikes (**b**) and effect of TTX on D-spike transitions (**c**). These data were not plotted in the main figures for sake of clarity. In **a**, **b** data are shown normalized to the average value below threshold (except for ∆F/F dendrite because of several points with value zero) and corrected for linear trend in subthreshold data (see Methods). In **c**, changes ∆ in parameter values across threshold in TTX are shown normalized to their increase ∆ in control, thus no correction for linear trends is required. Analysis of half duration is missing because there were not enough data points for statistical analysis.
